# Corticotropin-Releasing Hormone Signaling in the Oval Bed Nucleus of the Stria Terminalis Mediates Chronic Stress-Induced Negative Valence Behaviors Associated with Anxiety

**DOI:** 10.1101/572966

**Authors:** Pu Hu, Isabella Maita, Christopher Kwok, Edward Gu, Mark Gergues, Ji Liu, Zhiping Pang, Dick F. Swaab, Paul J. Lucassen, Troy A. Roepke, Benjamin A. Samuels

## Abstract

The bed nucleus of stria terminalis (BNST) is a forebrain region highly sensitive to stress that expresses corticotropin-releasing hormone (CRH) neuropeptide at high levels and regulates negative valence behaviors associated with anxiety. However, how chronic stress modulates CRH signaling and neuronal activity in BNST remains unknown. We subjected C57BL6/J mice to a 6-week chronic variable mild stress (CVMS) paradigm and investigated the effects on behavior, BNST cellular neurophysiology, and BNST CRH signaling. We also utilized pharmacological infusions and optogenetics to decipher and mimic the effects of CVMS on BNST cellular neurophysiology and behavior. CVMS elevated plasma corticosterone levels, induced negative valence behaviors associated with anxiety, diminished M-currents (voltage-gated K^+^ currents that stabilize membrane potential and regulate neuronal excitability), and increased mEPSC amplitude in ovBNST. Additionally, the number of c-fos^+^, CRH^+^, and CRH activator pituitary adenylate cyclase-activating polypeptide (PACAP)^+^ cells were increased while CRH inhibitor striatal-enriched protein tyrosine phosphatase (STEP)^+^ cells were decreased in ovBNST. These expression data were confirmed with qPCR. CVMS also activated PKA in BNST and the electrophysiological and behavioral effects of CVMS were reversed by ovBNST infusion of the PKA-selective antagonist H89. Moreover, optogenetic activation of ovBNST directly induced negative valence behaviors associated with anxiety, mimicking the effects of CVMS. CVMS mediates effects on negative valence behaviors associated with anxiety by activating CRH signaling components and cellular excitability in ovBNST Our findings decipher an important CRH-associated stress molecular signature in BNST and advance our understanding of the neural circuitry underlying stress-induced disorders.

## Introduction

Stressful environments promote vigilance, which, when moderate, is adaptive and essential for survival. However, chronic exposure to stressful experiences or environments can be maladaptive and increase risk assessment of low imminence threats, which is a feature of persistent anxiety. The limbic forebrain structure bed nucleus of the stria terminalis (BNST) is critical for mediating the neuroendocrine stress response (1) and potential threat regulation (2-5). Overall, BNST integrates stress and reward information from the limbic system and projects to neuroendocrine and autonomic neural systems located in the hypothalamus and brain stem regions that mediate the hypothalamic-pituitary-adrenal (HPA) stress response (3,5,6). Stimulating BNST elicits anxiogenic responses (7), whereas BNST inactivation is anxiolytic (8,9). Consequently, BNST dysfunction contributes to exaggerated stress responses and stress-related mood disorders.

The stress hormone corticotropin-releasing hormone (CRH or CRF) is abundantly expressed in hypothalamic paraventricular nucleus (PVN) parvocellular neurons. These PVN CRH neurons regulate HPA axis activity (10) and cortisol release (11-13). Importantly, CRH is also highly expressed in BNST (5, 14-18), especially in the oval nucleus (19). CRH mRNA in the antero-dorsolateral BNST (BNSTadl) is increased by stress exposure (20). Furthermore, anterolateral BNST (BNST_ALG_) stimulation increases corticosterone release (6,21). Notably, BNST orchestrates stress responses in a CRH-dependent fashion (22). CRH neurons in BNSTadl mediate anxiogenic effects (19,23,24) and negative affective responses to stress (25-28). CRH injections into BNST and CRH overexpression in BNST cause anxiogenic effects (26) and result in increased negative valence behaviors associated with anxiety (29,30). Therefore, CRH dysfunction in BNST likely contributes to stress-related behavioral states (31) and mood disorders (32).

BNST is a complex conglomerate structure with a heterogeneous population of neurons (19, 33-35). Within BNST_ALG_, the highest concentration of CRH neurons is located in the oval nucleus (14,19,36) (ovBNST). ovBNST is thought to be a master controller of BNST outflow that regulates overall BNST activity (3). Optogenetic inhibition of ovBNST deceases, whereas stimulation promotes negative valence behaviors (37). Here, we utilize a chronic variable mild stress (CVMS; also known as chronic unpredictable mild stress) paradigm in mice to better understand the effects of chronic stress on BNST CVMS and other chronic stress paradigms are widely used to determine the neuroendocrine and physiological effects that result in behavioral disturbances in stress-related mood disorders (38) (39). However, little is known about how chronic stress modulates the expression of CRH signaling components and cellular neurophysiology properties in the BNST and whether these alterations underlie chronic stress effects on behavior. Here we demonstrate that CVMS modulates expression of CRH-related stress signaling components (including its upstream activator, pituitary adenylate cyclase-activating polypeptide (PACAP) as well as its inhibitor, striatal-enriched protein tyrosine phosphatase (STEP)) and neuronal activity as measured by both M-currents (a voltage-dependent non-inactivating K^+^ current that stabilizes cellular membrane potential) and miniature excitatory postsynaptic currents (mEPSCs) in mouse ovBNST. Furthermore, we show that selective inhibition of PKA (a downstream modulator of CRH signaling) in ovBNST reverses CVMS effects on M-currents, mEPSCs, and behavior. Furthermore, optogenetic activation of ovBNST mimics CVMS effects on behavior. Taken together, these data suggest that the effects of CVMS on negative valence behaviors associated with anxiety are mediated through altered CRH signaling and neuronal activity in ovBNST.

## METHODS AND MATERIALS

### Mice

All procedures were in accordance with institutional guidelines based on the National Institutes of Health standards and approved by the Rutgers Institutional Animal Care and Use Committee. All mice used (except for optogenetics) were adult male C57BL/6J mice purchased from Jackson. For optogenetics, dopamine receptor D1a (Drd1a)-Cre (GENSAT line EY266) transgenic mice were generated from in house breeding.

Except for CVMS exposure, all mice were maintained in a controlled temperature (22°C) and a 12 h light/dark cycle with food and water provided *ad libitum.*

### Chronic variable mild stress (CVMS)

CVMS was performed as described (40,41). CVMS began when mice were 6 weeks old and persisted for 6 weeks. 50 mice were randomly assigned to non-stress (Control) (n=25) or CVMS groups (n=25). Variable mild stressors were used: daily bedding alterations (repeated sawdust changes, removal of sawdust, damp sawdust, substitution of sawdust with 21 °C water), cage-tilting (45° angle), predator sounds (15 min), cage shift (placed into the empty cage of another male), alterations of the light/dark cycle, lights off for 180 min, overnight food/water deprivation (40,41).

50 total mice were used in 3 sets of experiments. 20 were used for behavior (n=10 per group) and subsequent immunohistochemistry (IHC) (randomly selected n=6 per group), 12 for electrophysiology (n=6 per group), and 18 for plasma CORT measurements (n=7-9 per group) and real-time quantitative PCR analysis (q-PCR) (n=7-9 per group).

### Data Analysis

All data are presented as mean±SEM. Statistical analysis were conducted with GraphPad Prism (La Jolla, CA). Comparisons of M-current *I-V* plots between Control and Stress group were performed at each voltage (−25 to −75 mV) using a one-way ANOVA with *posthoc* Newman-Keuls comparisons. Maximum current at −35 mV was analyzed with paired Student’s t-test. For mEPSCs, amplitude and frequency were analyzed off-line using Mini Analysis (Synaptosoft, Fort Lee, NJ). mEPSC amplitude and frequency comparisons were performed using paired Student’s *t*-test. For behavior, IHC, and plasma CORT concentration comparisons between Control and CVMS groups, data were analyzed with a one-way ANOVA and *posthoc* Tukey comparisons. For optogenetics behavior experiments, data were analyzed using a two-way ANOVA with *posthoc* Tukey comparisons, with light and group as independent factors.

Differences were considered significant when p<0.05. *n* represents the number of cells or animals.

Additional methods and materials are provided in the Supplemental File.

## RESULTS

We first exposed a cohort of male C57BL6/J mice to either chronic variable mild stress (CVMS) or control conditions. CVMS began when mice were 6 weeks old and the paradigm persisted for 6 weeks (Fig.1A). 6 weeks of CVMS resulted in a significant percent decrease in body weight compared with non-stress controls (F(1,13)=84.908, p<0.001). We next sampled blood plasma and found significantly higher basal CORT levels in the CVMS group relative to the control group (F(1,18)=12.281, p<0.01) (Fig.S1).

**Fig.1:**
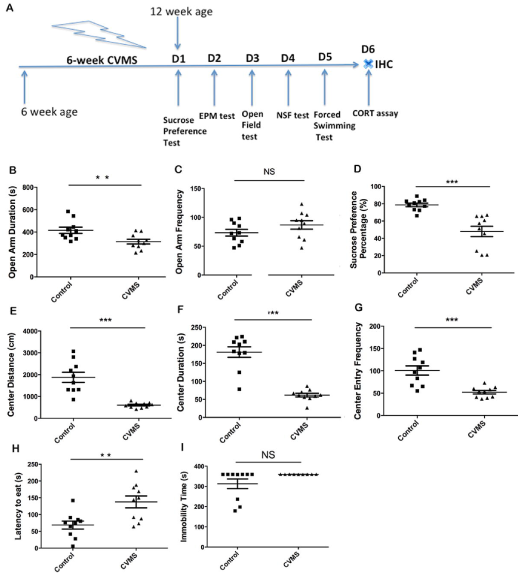
Schematic for experimental design and results for a series of negative valence behavior tests after mice were subjected to chronic variable mild stress (CVMS) paradigm. (A): Mice were subjected to a 6-week chronic variable mild stress (CVMS) paradigm, then subjected to a series of anxiety/depressive-like behavioral tests including sucrose preference test, elevated plus maze (EPM) test, open field (OF) test, novelty suppression feeding (NSF) test and forced swimming test, then were perfused for immunohistochemistiy (IHC) study. (B): Duration in open arm compared between control (n=10) vs. CVMS (n=10) mice in the elevated plus maze (EPM) test. (C): Frequency in the open arm compared between control (n=10) vs. CVMS (n=10) mice in the elevated plus maze (EPM) test. (D): Sucrose preference percentage compared between control (n=10) vs. CVMS (n=10) mice. (E): The distance that the mice travelled in the center of open field (OF) test compared between control (n=10) vs. CVMS (n=10) mice. (F): The duration that the mice spent in the center of open field (OF) test compared between control (n=10) vs. CVMS (n=10) mice. (G): The frequency that the mice entried into the center of open field (OF) test compared between control (n=10) vs. CVMS (n=10) mice. (H): Comparison of the latency to eat food pellets in the novelty suppressed feeding (NSF) test between control (n=10) vs. CVMS (n=10) mice. (I): Comparison of the immobility time in the forced swim test (FST) between control (n=10) vs. CVMS (n=9) mice. n=9-10 animals per group; **: p<0.01; ***: p<0.001.

We next assessed behaviors that are influenced by chronic stress exposure. Mice exposed to CVMS showed decreased open arm duration in elevated plus maze (EPM) (F(1,18)=8.262; p=0.01; Fig.1B), decreased sucrose preference in sucrose preference test (SPT) (F(1,18)=23.118; p<0.001; Fig.1D), decreased center distance (F(1,18)=30.155; p<0.001; Fig.1E), center duration (F(1,18)=59.24; p<0.001; Fig.1F), and center entries (F(1,18)=19.50; p=0.001; Fig.1G) in open field (OF), and increased latency to eat in novelty suppressed feeding (NSF) (F(1,18)=10.460; p<0.01; Fig.1H) relative to non-stress controls. EPM open arm entries (F(1,18)=2.065; p=0.17; Fig.1C) and forced swim test (FST) immobility (F(1,18)=3.035; p=0.08; Fig.1I) were not significantly affected by CVMS. Taken together, these data demonstrate that our CVMS protocol effectively induces negative valence behaviors associated with anxiety in EPM, OF, and NSF, and decreases reward valuation in the anhedonia-related SPT

Next, we determined the effects of CVMS on ovBNST electrophysiological properties using *ex vivo* slices (Fig.2A). We first measured M-currents (KCNQ/Kv7 channels), a subthreshold noninactivating voltage-dependent outward K^+^ current that controls action potential generation and neuronal excitability (42), in ovBNST neurons using a standard deactivation-activation protocol over a voltage range (−75 to −25 mV) where M-currents have profound effects on neuronal excitability (Fig.2C). M-currents were calculated by determining current relaxation, the difference between the instantaneous and steady states (Fig.3C arrows). The maximum M-currents were recorded at −35 mV (Fig.2B). 20 min of recording did not show a M-current rundown (amplitude decrease) (example traces in Fig.2B; n=5). Application of the KCNQ/Kv7 channel blocker XE991 robustly decreased M-currents (Fig.2B). We next determined the role of M-currents in modulating BNST neuronal excitability. To this end, firing activity of ovBNST neurons was continuously monitored in current clamp mode (Fig.2D). XE991 application induced action potentials after 30 s, and 6-7 min of XE991 perfusion led to robust firing bursts indicative of activation and hyperexcitability (Fig.2D).

**Fig.2:**
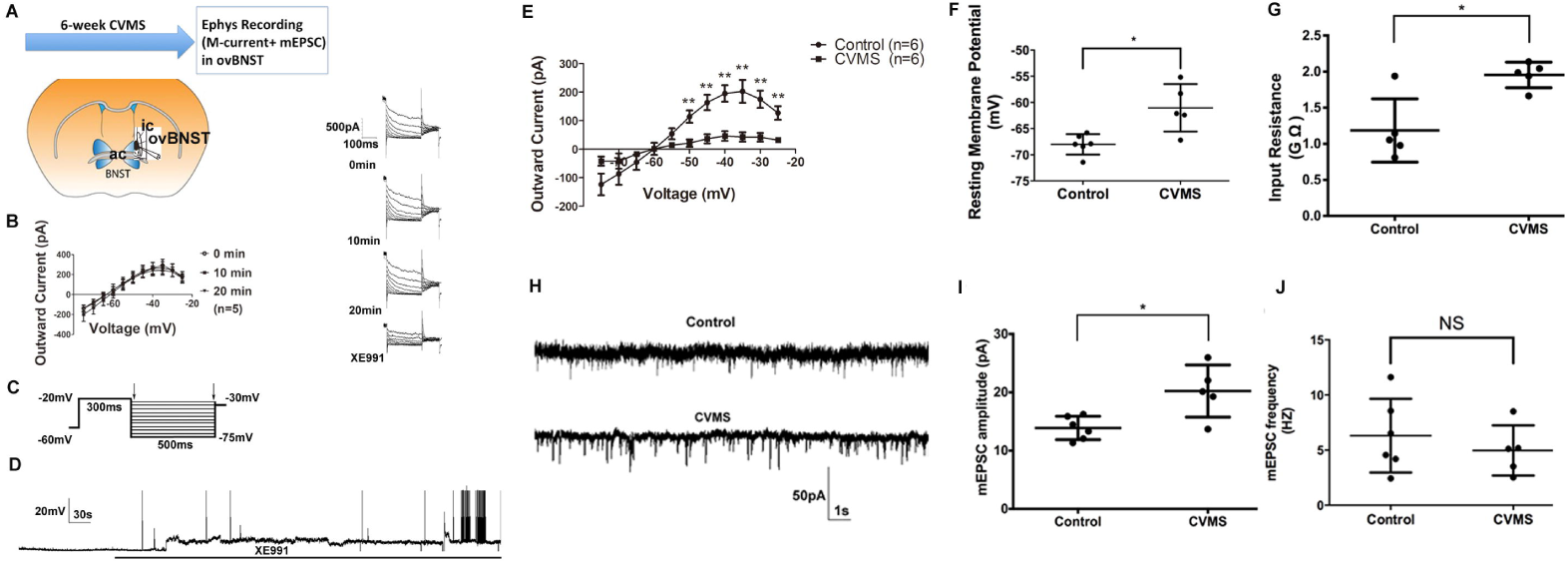
Anatomy example of mice oval nucleus of the bed nucleus of the stria terminalis (ovBNST) and chronic variable mild stress (CVMS) effects on the electrophysiological properties (including M-current recording and miniature excitatory postsynaptic current (mEPSC)) in the mice ovBNST. (A): Anatomy example of ephys recording site in the oval nucleus of mice bed nucleus of the stria terminalis (ovBNST) by whole-cell patch clamping method; ic: internal capsule; ac: anterior commissure. (B): Left: I-V plot of M-current from −75 mV to −25 mV shows that M-current recorded in ovBNST did not run down after 20 min; Right: example of M-current traces shown at 0 min, 10 min and 20 min and after subsequent perfusion with KCNQ-selective channel blocker XE991 (10 min, 40 μM). (C): The deactivation protocol used to record the M-current: from a holding potential of −60 mV, a voltage jump to −20 mV (300 ms) was followed by steps from −30 mV to −75 mV in 5 mV increments (500 ms); (D): An example of continuous action potential firing activity in a recorded ovBNST neuron during perfusion of selective KCNQ channel blocker XE991 (40 μM) for 10 min after 2 min baseline recording under current-clamp mode. Robust action potential is seen starting 7min after XE991 perfusion. (E): I-V plot shows significantly diminished outward M-current in ovBNST neurons ranging from −75 mV to −25 mV compared between control vs. CVMS mice (both n=6 cells). Significant difference was found between the voltage range of −50 to −25 mV. (F): Comparison of cellular resting membrane potential (RMP) in ovBNST neurons from control (n=6 cells) vs. CVMS mice (n=5 cells) revealed significant higher RMP in the ovBNST of CVMS mice. (G): Comparison of cellular input resistance (IR) in ovBNST neurons from control (n=5 cells) vs. CVMS mice (n=5 cells) revealed significant higher IR in the ovBNST of CVMS mice. (H): Example of a comparison of mEPSC traces in ovBNST from control vs. CVMS mice. CD: Average mEPSC amplitude increased in ovBNST of CVMS mice (n=6 cells) compared to Control mice (n=5 cells). (J): Average mEPSC frequency did not change in ovBNST of CVMS mice (n=6 cells) compared with Control mice (n=5 cells). *: p<0.05; **: p<0.01; NS: non-significant different (p>0.05).

Interestingly the outward M-current was attenuated in ovBNST neurons from CVMS mice (Fig.2E), especially at higher voltages (p=0.005, 0.003, 0.001, 0.004, 0.003, and 0.005 at −50, −45, −40, −35, −30 and −25 mV, respectively; n=6), with a significant effect of stress (F(1,10)=16.353, p=0.002). At −35 mV, the outward M-current peak value was robustly decreased from 202.59±39.95 pA in control to 42.90±16.54 pA (t=3.693; p=0.004) in CVMS slices. Furthermore, CVMS significantly depolarized the resting membrane potential (RMP) (t=3.192, p=0.023) (Fig.2F) and increased the input resistance (Rin) (t=3.632, p=0.014) (Fig.2G) in ovBNST neurons.

Reduced M-currents can elicit increased excitatory cellular responses to synaptic inputs (43). Therefore, we hypothesized that CVMS may alter excitatory glutamatergic neurotransmission in ovBNST Indeed, CVMS significantly increased the average amplitude of mEPSCs (Fig.2H-J) (t=3.141, p=0.012) without affecting mEPSC frequency (t=0.790, p=0.451 (Fig.2J). Taken together, these data demonstrate that CVMS contributes to increased excitability and hyperactivation of ovBNST neurons.

Corticotropin-releasing hormone (CRH), a neuropeptide that is released from the paraventricular nucleus (PVN) of the hypothalamus (PVN) in response to stress, is also highly expressed in BNST (Fig.3.1A). Therefore, we next assessed CVMS effects on CRH signaling in BNST. To this end, we subdivided anterior-dorsolateral BNST (BNSTadl) into the oval nucleus of BNST (ovBNST) and the surrounding anterolateral dorsal region of BNST (adBNST) (Fig.S4E). CVMS increased the number of CRH^+^ cells in BNSTadl and ovBNST relative to control (BNSTadl: F(1,1G)=49.78, p<0.01; ovBNST: F(1,10)=28.93; p<0.01) (Fig.3.2C-D). However, the number of CRH^+^ cells in adBNST did not differ between CVMS and control (Fig.S4.2A).

**Fig.3_1:**
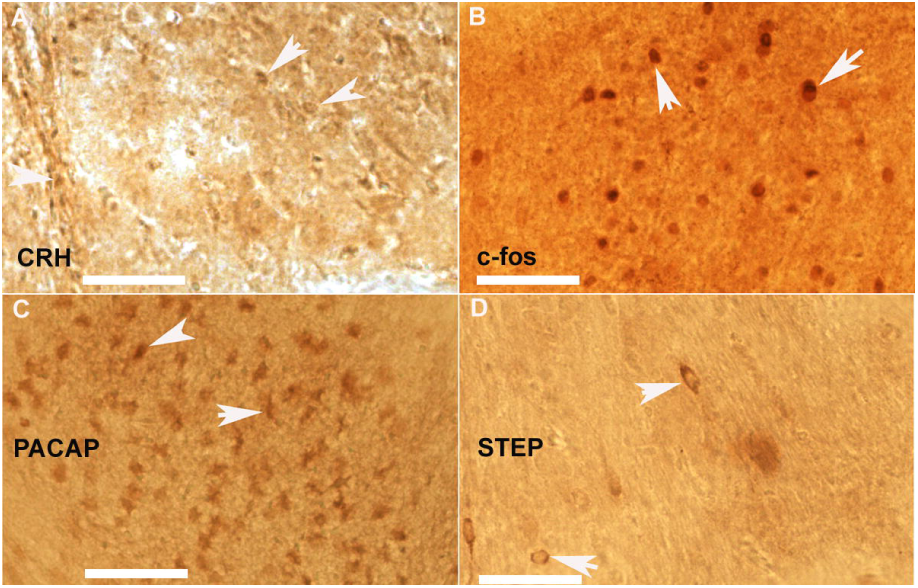
Typical example of high magnification figure showing immunostaining of CRH (A), c-fos (B), PACAP (C) and STEP (D) in the mice oval nucleus of BNST (ovBNST). (A): White arrows point to typical CRH-immunoreactive (IR) cells in the ovBNST: (B): White arrows point to typical c-fos-immunoreactive (IR) cells in the ovBNST; (C): White arrows point to typical PACAP-immunoreactive (IR) cells in the ovBNST: (D): White arrows point to typical STEP-immunoreactive (IR) cells in the ovBNST: All figures are in 40X magnification. Scale bar: 50iim. ic: internal capsule; ac: anterior commissure.

**Fig.3_2:**
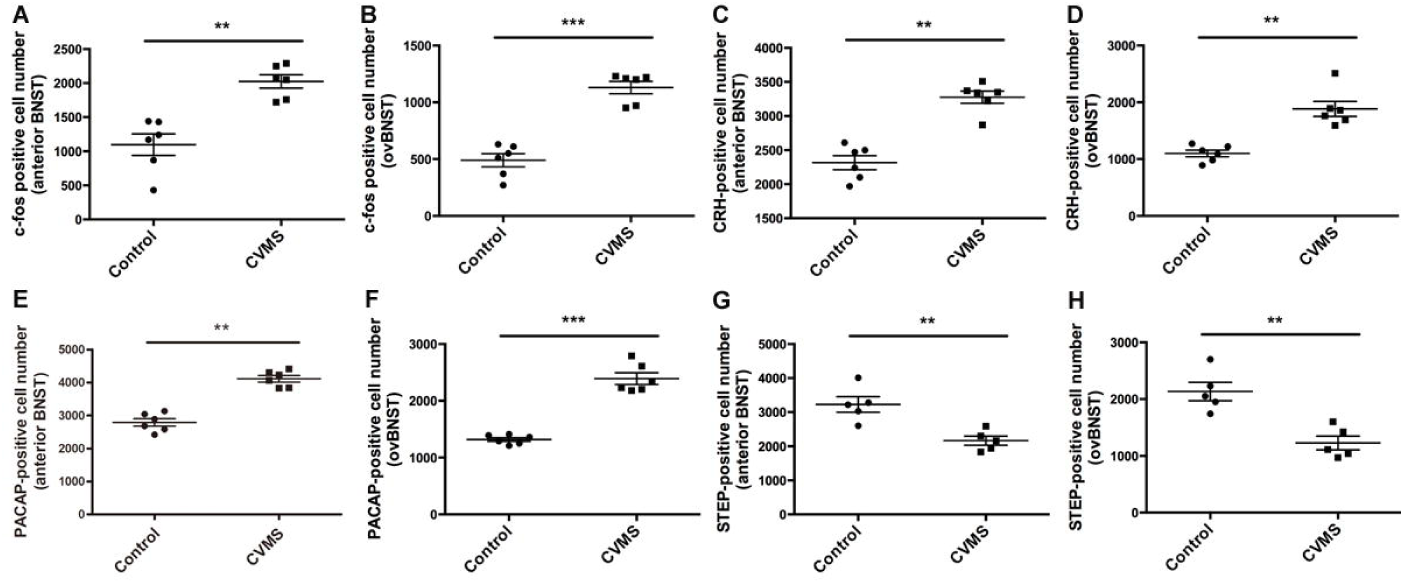
Comparison of c-fos (A,B), CRH (C,D), PACAP (E,F) and STEP (G,H)-immunoreactive (IR) cell numbers in both whole antero-dorsolateral BNST (BNSTadl) and the oval nuclei of BNST (ovBNST). (A) and (B): Comparison of c-fos-IR cell number in the (A) BNST_adl_ and (B) ovBNST shows higher number of c-fos-IR cells in both regions of BNST from CVMS mice (n=6) compared to Control mice (n=6);.. (C) and (D): Comparison of CRH-IR cell number in the (C) BNST_adl_ and (D) ovBNST shows higher number of CRH-IR cells in both regions of BNST from CVMS mice (n=6) compared to Control mice (n=6);. (E) and (F): Similarly, comparison of PACAP-IR cell number in the (E) BNST_adl_ and (F) ovBNST shows higher number of PACAP-IR cells in both regions of BNST from CVMS mice (n=6) compared to Control mice (n=6); (G) and (H): On the contrary comparison of STEP-IR cell number in the (G) BNST_adl_ and (H) ovBNST reveals lower number of STEP-IR cells in both regions of BNST from CVMS mice (n=6) compared to Control mice (n=6). **: p<0.01; ***:p<0.001

We next assessed c-fos in the BNST of control and CVMS mice as a marker of neuronal activation (c∼fos+ cell example in Fig.3.1B). The number of c-fos-immunoreactive cells in BNSTadl and ovBNST were significantly increased by CVMS (BNSTadl: F(1,10)=24.896, p<0.01; ovBNST: F(1,10)=65.29, p<0.001) (Fig.3.2A-B). However, c-fos^+^ cells were unchanged by CVMS in the adBNST region (Fig.S4.2B).

Consistent with these data, the CRH activator pituitary adenylate cyclase-activating polypeptide (PACAP) (Fig.3.1C), was more highly expressed in BNSTadl and ovBNST after CVMS, while the CRH inhibitor striatal-enriched protein tyrosine phosphatase (STEP) (Fig.3.1D), was decreased in BNSTadl and ovBNST of CVMS mice (PACAP BNSTadl: F(1,10)=77.89, p<0.01; PACAP ovBNST: F(1,10)=98.36, p<0.001 (shown in Fig.3.2.E and F)); STEP BNSTadl: F(1,8)=16.063, p<0.01; STEP ovBNST: F(1,8)=20.099, p<0.01) (Fig.3.2G-H). However, the number of PACAP^+^ and STEP^+^ cells in adBNST was unchanged by CVMS (Fig.S4C-D). Taken together, these data demonstrate that CRH signaling is increased in BNSTadl and ovBNST by CVMS.

To complement these immunohistochemistry results, we next assessed mRNA expression of CRH, PACAP, STEP, and the CRH receptors CRHR1 and CRHR2 in BNSTadl by qPCR (Fig.4A). CVMS significantly increased CRH (F(1,14)=6.303, p=0.018; Fig.4B) and PACAP (F(1,12)=5.597, p=0.037; Fig.4C) and decreased STEP (F(1,16)=5.877, p=0.031; Fig.4D) expression in BNSTadl relative to non-stress controls. Interestingly, CRHR1 expression was also significantly increased in CVMS mice (F(1,15)=4.985, p=0.041; Fig.4E), whereas CRHR2 was unchanged (F(1,14)=0.750, p=0.403; Fig.4F). CVMS did not affect KCNQ2, KCNQ3, and KCNQ5 subunit (the Kv7 M-channel components) expression (Fig.S2). Since these qPCR results suggest that CRHR1 expression is selectively increased in the BNST by CVMS, and CRHR1 is a Gs protein-coupled receptor linked to adenylyl cyclase (AC), we next assessed whether CVMS resulted in protein kinase A (PKA) activation in BNST. To this end, we assessed protein expression levels of PKA (using an antibody recognizing the PKA subunit C-α) and its activated form p-PKA (using an antibody recognizing Thr197 phosphorylated-PKA-C; Fig.4G) in BNSTadl tissue punches from CVMS and control mice by western blot. While total PKA levels did not differ (t=1.46, p=0.171) (Fig.4l), p-PKA expression in BNSTadl was increased by CVMS (*t* test t=5.105, p=0.0003) (Fig.4J), indicating CVMS increases PKA activation in BNSTadl.

**Fig.4:**
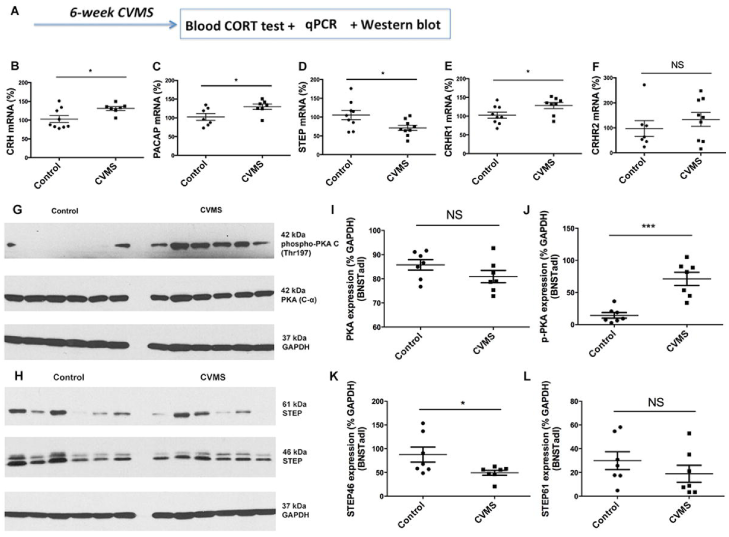
Schematic experimental design and results for qPCR and western blot study after mice were subjected to 6-week chronic variable mild stress (CVMS) paradigm. (A): Experimental design for qPCR and western blot study to detect mRNA and protein expression changes in the antero-dorsal lateral region of the BNST (BNST_adl_) after 6-week CVMS paradigm. (B): Higher expression of CRH mRNA was found in BNSTadl of CVMS mice (n=7) vs. control mice (n=9); (C): Similarly PACAP mRNA expression was higher in BNSTadl from CVMS (n=7) mice vs. control (n=7) mice; (D): On the contrary comparison of STEP mRNA expression in BNSTadl between control (n=9) vs. CVMS (n=9) mice revealed lower STEP mRNA expression from CVMS mice; (E): CRHR1 mRNA expression in BNSTadl of CVMS (n=8) mice is higher compared to control (n=9) mice; (F): No significant change was found for the mRNA expression of CRHR2 in BNSTadl compared between control (n=7) vs. CVMS (n=9) mice; (G): Example of western blot example showing comparison of 42 kDa phospho-PKA C (Thrl97) (1^st^ lane) and 42 kDa PKA (C-α) (2^nd^ lane) in BNSTadl from control (Left; n=6) vs. CVMS (Right; n=6) mice; 37 kDa GAPDH was used as endogenous control; (H): Example of western blot example showing comparison of expression of 61 kDa membrane isoform of STEP61 and 46 kDa cytosolic isoform of STEP46 in BNSTadl from control (n=6) vs. CVMS (n=6) mice; 37 kDa GAPDH was used as endogenous control; (I): Quantification of PKA protein expression level in BNSTadl between control (n=6) vs. CVMS (n=6) mice; (J): Quantification and comparison of p-PKA (Thrl97) expression level in BNSTadl from control (n=6) vs. CVMS (n=6) mice revealed higher expression of p-PKA from the CVMS mice compared to Control mice; (K): Quantification of comparison of cytosolic STEP46 revealed higher expression level in the BNSTadl from control (n=6) vs. CVMS (n=6) mice; (L): Quantification of comparison of membrane STEP61 expression level in BNSTadl from control (n=6) vs. CVMS (n=6) mice revealed no significance change compared between these 2 groups. *: p<0.05; ***: p<0.001; NS: non-significant different (p>0.05).

We also assessed STEP protein expression in BNSTadl tissue punches (Fig.4H). Whereas the membrane isoform STEP61 did not differ (t=1.060, p=0.31) (Fig.4L), CVMS decreased expression of the cytosolic isoform STEP46 in BNSTadl was decreased in CVMS mice (t=2.292, p=0.026) (Fig.4K), suggesting CVMS specifically affects cytosolic STEP46 expression in BNST.

We next assessed whether BNST PKA activation mediates the effects of CVMS on M-currents and mEPSCs. When *ex vivo* brain slices were pre-incubated with the selective PKA antagonist H89 for 30 min, the CVMS-induced decrease in ovBNST outward M-currents was significantly attenuated at higher voltages (F(1,10)=16.353, p=0.002; Fig.5A) (p=0.044, 0.045, 0.047, 0.042 and 0.020 at −45, −40, −35, −30 and −25 mV, respectively). At −40 mV, the outward M-current peak value was increased (t=2.292, p=0.045) in CVMS slices pre-incubated with H89 compared with CVMS slices (Fig.5A). Similarly, the CVMS-induced mEPSC amplitude increase in ovBNST was significantly decreased (t=2.913, p=0.025) when slices were preincubated with the selective PKA antagonist H89 for 30 min (Fig.5B). By contrast, mEPSC frequency in ovBNST was not different between CVMS and CVMS+H89 slices (t=0.210; p=0.843) (Fig.5C). Importantly, H89 pre-incubation had no significant effects on ovBNST M-currents and mEPSCs in control slices (Fig.S8). Taken together, these results suggest that the effects of CMVS on M-currents and mEPSC amplitude in ovBNST are mediated by PKA activation.

**Fig.5:**
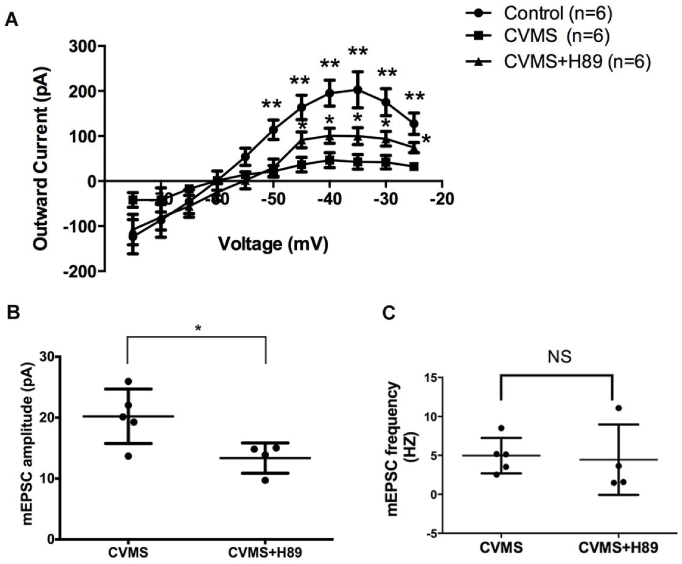
Effects of CVMS on the M-currents and mEPSC recorded in the ovBNST are partly mediated by PKA-activation. (A): BNST brain slices from CVMS mice were pre-incubated with PKA-selective antagonist H89 for 30 min to compare its effect on the properties of M-current and mEPSC recorded in the oval nucleus of BNST (ovBNST) of the chronic stress (CVMS) mice. I-V curve of outward M-current shows that the attenuated M-currents in ovBNST of CVMS mice were partly normalized at the membrane voltage ranging from −45 mV to −25 mV after slices were pre-incubated with PKA-selective antagonist H89 (n=6 cells). (B): H89 (n=6 cells) pre-incubation normalized the increased mEPSC amplitude in ovBNST of CVMS mice (n=5 cells) compared with Control mice. (C): H89 (n=6 cells) pre-incubation had no significant effect on the mEPSC frequency in ovBNST of CVMS mice (n=5 cells). *: p<0.05; ***: p<0.001; NS: non-significant different (p>0.05).

Next, we examined whether PKA activation also mediates the behavioral effects of CVMS by infusing the PKA-selective antagonist H89 (25 nM, dissolved in 0.5 μl saline) into ovBNST (Fig.6A-B). BNST H89 infusions into CVMS mice significantly increased EPM open arm duration (F(1,14)=12.528; p=0.011; Fig.6C), OF center distance (F(1,15)=27.962; p<0.01; Fig.6E), OF center duration (F(1,15)=31.752; p<0.01; Fig.6F), OF center entries (F(1,15)=18.121; p<0.01; Fig.6G), and SPT sucrose preference (F(1,19)=4.43; p=0.035; Fig.6H), and decreased NSF latency to eat (F(1,19)=4.434; p=0.025) (Fig.6I) relative to control infusions into CVMS mice. H89 did not affect EPM open arm entries in CVMS mice (F(1,14)=1.212; p=0.374; Fig.6D). Importantly, H89 infusions into BNST had no effects in EPM, SPT, OF, or NSF in control mice (Fig.S9). Taken together, these results demonstrate that CVMS effects on negative valence behaviors associated with anxiety in tasks such as EPM, OF, and NSF and reward valuation in the anhedonia-related SPT are mediated by ovBNST PKA activation.

**Fig.6:**
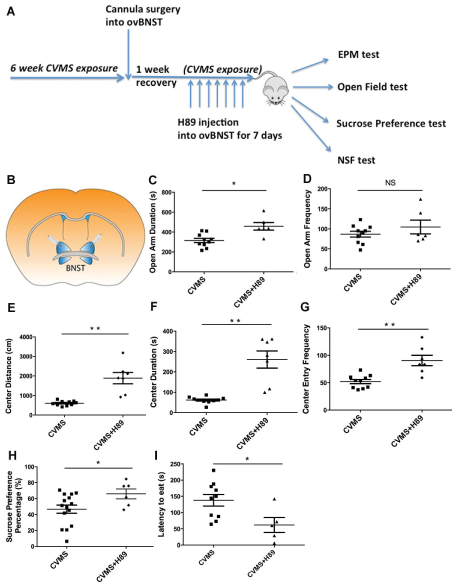
PKA activation in ovBNST mediates CVMS effects on negative valence behaviors associated with anxiety. (A): Cannula surgery schematic showing that PKA-selective antagonist H89 was chronically infused into the oval nucleus of BNST (ovBNST) of CVMS mice for 7 days continually to assess behavioral effects of CVMS in EPM, OF, SPT, and NSF test). CVMS exposure was continuously present during the chronic 7-day drug infusion period. Mice was allowed to recover for 7 days before chronic H89 injection starts after cannula surgery. (B): Anatomy example shows cannula inserted into ovBNST. (C): PKA-selective antagonist H89 (Stress+H89; n=7) significantly increased the duration time that mice spent in open arm of EPM test compared with CVMS mice (Stress; n=10). (D): H89 (Stress+H89; n=7) had no significant effect on the frequency that mice entried into open arm of EPM test compared with CVMS mice (Stress; n=10). (E): H89 (Stress+H89; n=7) significantly increased the distance that the CVMS mice (Stress; n=10) traveled in the center of open field (OF) test. (F): H89 (Stress+H89; n=7) significantly increased the duration that the CVMS mice (Stress; n=10) spent in the center of open field (OF) test. (G): H89 (Stress+H89; n=7) significantly increased the frequency that the CVMS mice (Stress; n=10) entried in the center of open field (OF) test. (H): H89 (Stress+H89; n=7) significantly increased the sucrose preference percentage of CVMS mice (Stress; n=10). (I): H89 (Stress+H89; n=7) significantly decreased the latency that the CVMS mice (Stress; n=10) start to eat food pellet of novelty suppressed feeding (NSF) test.

Since our data demonstrate that CVMS decreases M-currents and increases mEPSC amplitude in BNST, we next sought to determine whether selective activation of ovBNST mimics the behavioral effects of CVMS. To this end, we injected either a Cre-inducible ChR2 (AAV5-EF1a-DIO-ChR2(H134R)-eYFP) or control virus (AAV5-EF1a-DIO-eYFP) into BNST of dopamine receptor D1a (Drd1a)-Cre transgenic mice (Fig.7A). Within BNST, Cre expression in Drd1a-Cre mice is restricted to ovBNST (37). Fig.7B-C shows expression of Control-eYFP (B) and ChR2-eYFP virus (C) in ovBNST. Similar to a previous report (Kim et al 2013), ChR2 photoillumination in ovBNST resulted in a significant virus *x* light interaction for EPM open arm duration and entries (Fig.7D-E), (Duration: F(2,42)=3.277, p=0.048; Entries: F(2,48)=8.746, p=0.001). Specifically, ovBNST activation (light ON period) decreased open arm duration and entries relative to the preceding (Duration: p<0.01; Fig.7D; Entries: p<0.01; Fig.7E) and succeeding (Duration: p=0.018; Entries: p<0.01) light OFF periods. By contrast, there were no significant changes in open arm duration or entries across the light OFF/ON/OFF periods in mice injected with control virus. Similarly, in OF, ChR2 photoillumination in ovBNST resulted in a significant virus *x* light interaction for center distance (Fig.7F), center duration (Fig.7G), and center entries (Fig.7H) (Distance: F(2,39)=6.853, p=0.003; Duration: F(2,39)=4.72, p=0.015; Entries: F(2,36)=11.495, p<0.001). Specifically, ovBNST activation decreased these OF center measures relative to the preceding (p<0.01 for all center measures) and succeeding (p<0.01 for all center measures) light OFF periods (Fig.7F-H). By contrast, there were no changes in center measures across the light OFF/ON/OFF periods in mice injected with control virus. Taken together, these data demonstrate that selective activation of ovBNST neurons mimics the behavioral effects of CVMS and that ovBNST activation is sufficient to induce negative valence behaviors.

**Fig.7:**
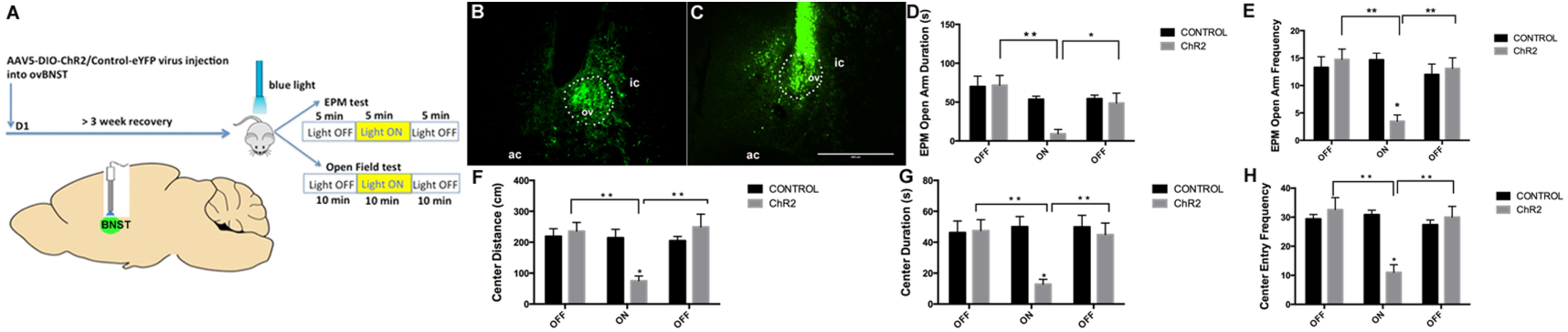
Optogeneticallv stimulating oval nucleus in the BNST (ovBNST) promotes negative valence behaviors associated with anxiety in EPM and OF. (A): Schematic graph showing the timeline of opsin virus (AAV5-EFlα-DIO-ChR2(H134R)-eYFP, ChR2) or control virus (AAV5-EFlα-DIO-eYFP, Control) injected into ovBNST of dopamine receptor D1a (Drd1a)-Cre transgenic mice. Mice were allowed to undergo at least 3 weeks recoveiy before subjected to either EPM or OF test (both composed of 3 different sessions of lights off-on-off series) under blue light stimulation. (B) and (C) shows the expression of Control-eYFP virus (B) and ChR2-eYFP virus (C) in ovBNST. Scale bar: 400 |im. Ic: internal capsule; ac: anterior commissure; ov: oval nucleus. (D): Comparison of the duration time that Control (n=8) mice vs. ChR2 (n=8) mice spent in open arm during the OFF-ON-OFF session (each 5 min) in EPM test revealed decreased open arm duration when blue light on only in the ChR2 group but not in the Control group. (E): Comparison of the frequency that Control (n=8) mice vs. ChR2 (n=8) mice entered into open arm during the OFF-ON-OFF session (each 5 min) in EPM test revealed decreased open arm frequency when blue light on only in the ChR2 group but not in the Control group. (F): Comparison of the distance that Control (n=7) mice vs. ChR2 (n=8) mice traveled in the center area during the OFF-ON-OFF session (each 10 min) in open Field test revealed decreased traveling distance in the OF center when blue light on only in the ChR2 group but not in the Control group. (G): Comparison of the time duration that Control (n=7) mice vs. ChR2 (n=8) mice spent in the center area during the OFF-ON-OFF session (each 10 min) in open Field test revealed decreased duration spent in the OF center when blue light on only in the ChR2 group but not in the Control group. (H): Comparison of the frequency that Control (n=7) mice vs. ChR2 (n=8) mice entried into the center area during the OFF-ON-OFF session (each 10 min) in open Field test revealed decreased entry frequency into the OF center when blue light on only in the ChR2 group but not in the Control group. *: p<0.05; **: p<0.01.

## DISCUSSION

Chronic stress results in long-term alterations in HPA axis activity and negative valence behaviors associated with anxiety (44-47). However, the neurobiological underpinnings remain elusive. BNST plays an important role in mediating sustained negative valence behaviors in rodents (8,48,49) and mood disorders in humans (50-54), and BNST dysfunction is implicated in stress-related psychopathology (2-4). Importantly, our data demonstrate that CVMS alters expression of CRH-related stress signaling components (including PACAP, STEP, and CRHR1) and enhances cellular excitability (manifested by decreased M-current and increased mEPSC amplitude) in ovBNST neurons. These CVMS-induced alterations result in BNST dysfunction, which in turn contributes to negative valence behavioral disturbances associated with anxiety. Direct optogenetic activation of ovBNST neurons mimics behavioral effects of CVMS, further suggesting that BNST dysfunction may underlie CVMS-mediated behavioral disturbances.

The CVMS paradigm is well validated to elicit negative valence behaviors. Whereas chronic exposure to homotypic stressors usually engenders HPA habituation (55-58), this shortcoming is circumvented by the CVMS paradigm, in which rodents were exposed to randomly alternating stressors in an unpredictable manner (59). In this study, a robust behavioral phenotype and elevated plasma corticosterone levels were found after a 6-week CVMS exposure. It will be informative for future studies to determine whether other chronic stress paradigms, such as chronic social defeat stress, elicit similar effects on ovBNST.

### M-current and mEPSC regulation by CVMS

The M-current is a subthreshold voltage-dependent non-inactivating outward K^+^ current that stabilizes membrane potential and sets cellular threshold for action potential firing (60,61). M-channels are composed of Kv7 subunits (KCNQ) (62), open at resting membrane potential, and undergo conformational changes that facilitate even more opening during depolarization, which results in membrane potential clamping. Therefore, M-currents function as a brake on repetitive action potential charges (62) and play an essential role in controlling neuronal excitability (63). Diminished M-currents allow neurons to fire more rapidly (61), as evidenced during XE991 (M-channel selective blocker) perfusion. Our results show that CVMS-induced decreased M-currents in ovBNST are accompanied by a more depolarized cellular resting membrane potential (Fig.2F), indicating enhanced neuronal excitability. Acute restraint suppresses M-currents and increases cellular activity in CRH neurons in the hypothalamic PVN, which may contribute to HPA hyperactivation after acute stress (64). However, whether chronic stress modulates M-currents was previously unknown. Our study provides a direct cellular mechanism by which CVMS increases excitability of ovBNST neurons. Consequently, targeting ovBNST M-channels may provide a potential therapeutic strategy to treat stress-related mood disorders (65).

Our qPCR results revealed no significant changes in KCNQ subunit mRNA expression after CVMS (Fig.S2), so post-translational modifications such as phosphorylation may account for M-current inhibition. We found that CVMS effects on M-currents are mediated in part through post-translational PKA activation (Fig.4J). Although few studies have assessed direct regulation of KCNQ channel activity by PKA, M-currents can be inhibited by phosphorylation (15). Anchoring proteins, such as members of the A-kinase anchoring protein family (AKAP), may bind to and localize activated PKA into a membrane signaling complex near the KCNQ channel subunit in order to modulate channel function (17) (66). Similarly, increased amplitude (but not frequency) of mEPSCs after CVMS suggests a postsynaptic effect, such as PKA-mediated phosphorylation of the GluR1 subunit of AMPA receptors (24,67). GluR1 phosphorylation is closely correlated with increased receptor surface trafficking and membrane redistribution (68,69), and increased AMPA receptor density on postsynaptic membranes would increase mEPSC amplitude (reflecting increased synaptic responses) (70-72).

### CVMS-induced alterations in BNST CRH signaling

Although hypothalamic CRH signaling was not assessed here, previous studies show PVN CRH neuron activation (73) and increased PVN CRH expression in rats exposed to CVMS (74). In BNST, CRH signaling is linked to negative affective states (25,26, 75-78). Increased CRH concentrations (79,80) and mRNA levels (20,81) are found in rat BNST after chronic stress, however whether chronic stress modulates CRH neuropeptides in mouse BNST was unknown. Notably, the oval nuclei in the BNST_ALG_, which resides dorsal to the anterior commissure (19), harbors the highest concentration of CRH neurons in BNST (14,36). Local CRH signaling in ovBNST is suggested to be stress responsive (28,35,82). Importantly, we also compared CRH expression in the anterodorsal BNST region (adBNST) surrounding ovBNST. Interestingly, CVMS selectively increases the number of CRH-immunopositive cells in ovBNST (with no changes in adBNST). In addition, CVMS selectively increases CRHR1 mRNA expression in BNSTadl, suggesting the effects of CRH are mediated mainly through this receptor. This is consistent with recent studies demonstrating stress effects on behavior depend on CRHR1 receptor activation (22) and are blocked by CRHR1 antagonist injections into BNST (25,26). CRHR1 and CRHR2 subtypes play opposing roles during the stress response (44,47), as CRHR1 activation initiates the stress response and results in negative valence behaviors (25,26, 83-85), whereas CRHR2 activation in the BNST mainly functions to terminate the stress response and facilitate stress recovery (86,87). Thus, exaggerated CRHR1 expression and a disruption in the expression balance of these two receptors may dampen stress-coping capability and precipitate stress-related psychopathology.

We also assessed the effects of CVMS on pituitary adenylate cyclase-activating polypeptide (PACAP) and striatal-enriched protein tyrosine phosphatase (STEP). PACAP is a key stress regulator (88-90) that is upstream of CRH and stimulates CRH production and secretion (91). PACAP dysregulation is implicated in depression (92) and PTSD in humans (93). PACAP is highly expressed in BNST and PACAP-containing neurons in ovBNST closely interact with CRH-containing neurons as part of the stress response. Chronic stress increases PACAP expression in the dorsal BNST_ALG_ (81,94), and PACAP infusion into BNST increases plasma corticosterone concentrations (95). Treatment with a PAC1 receptor antagonist attenuates behavioral and endocrine effects of stress (94). We found that increased CRH and CRHR1 levels were accompanied by increased PACAP expression in ovBNST following CVMS. We also investigated the effects of CVMS on STEP (also known as protein tyrosine phosphatase nonreceptor type 5, PTPN5). STEP is a brain-specific tyrosine phosphatase that dephosphorylates and inactivates several kinases (including ERK1/2 and p38MAPK, Fyn) (96) and STEP overexpression enhances stress resilience (97). Furthermore, STEP loss-of-function increases susceptibility to stress-induced cognitive deficits (98). Downregulation of STEP expression in BNST_ALG_ after chronic stress may contribute to a prolonged negative affective state (28). Importantly, in BNST, STEP is coexpressed in ovBNST CRH neurons and selectively buffers CRH neurons against overactivation after stress (28). While CVMS resulted in increased expression of CRH, CRHR1, and PACAP, STEP expression in BNST_ALG_ was decreased. These concomitant changes suggest that CVMS disrupts the delicate balance of CRH signaling and results in hyperactivation of CRH neurons in the BNST_ALG_-Together, our data show that CVMS causes disturbances in BNST CRH signaling that reflect a novel stress-associated molecular signature.

### Working Model

We propose that activation of CRHR1 in ovBNST by CRH stimulates Gs protein-coupled adenylyl cyclase (AC) activity (99), which results in cAMP generation and activation of cAMP-dependent PKA. PKA phosphorylates the KCNQ channel and the GluRl subunit of AMPA receptors, which mediate M-current inhibition (15) and increased mEPSC amplitude (100,101), respectively (Fig.8). CRH itself can also facilitate glutamatergic transmission in many brain regions (102-104), Increased mEPSC amplitude coupled with decreased M-currents contribute to hyperexcitation of ovBNST neurons following chronic stress. In turn, hyperactive ovBNST neurons disrupt neural circuitry and result in dysfunctional HPA axis activation through projections to PVN in the hypothalamus.

**Fig.8:**
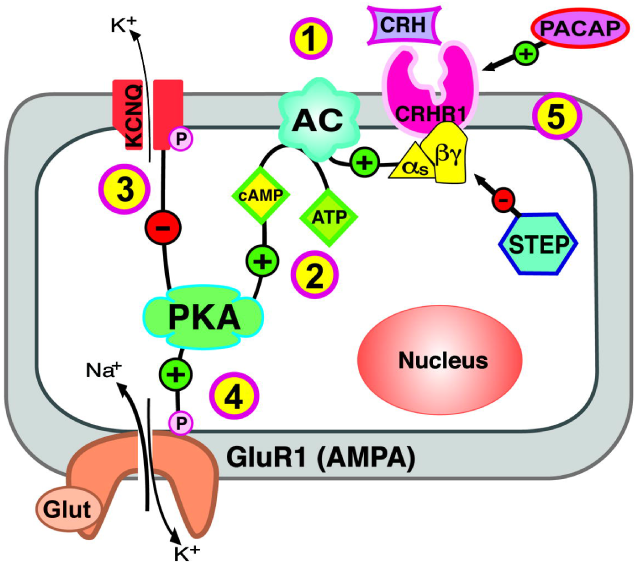
Working model illustrating the sequence of the molecular cascades of the CRH stress signaling in the BNST. Chronic stress 1): induces CRH production and release in the BNST, which sequentially activates CRHR1, a Gs-protein coupled membrane receptor. 1) Activation of AC (adenylyl cyclase) (coupled to Gs) then generates 2) cAMP production, which in turn 2): activates PKA enzyme. Activation of PKA then initiates two parallel phosphorylation pathways: 3) phosphorylates KCNQ channel on the cellular membrane to mediate inhibition of the M-current; and 4) phosphorylates GluRl subunit of AMPAR on the postsynaptic membrane to mediate potentiation of mEPSC amplitude. These two parallel pathways converge together to mediate M-current inhibition and increased mEPSC amplitude. Meanwhile: 5) PACAP (function as CRH activator) and STEP (function as CRH inhibitor) neuropeptide will function to activate and inhibit CRH respectively.

Acute optogenetic activation mimicked the effects of CVMS on ovBNST neurons. In these experiments, we utilized dopamine receptor D1a (Drd1a)-Cre transgenic mice where, within BNST, Cre is selectively expressed in ovBNST neurons (19,37,105). Similar to a previous report (Kim et al 2013), ChR2-mediated activation of ovBNST neurons induced negative valence behaviors associated with anxiety. Such acute effects on behavior are likely due to fast stimulatory release of local excitatory neurotransmitters, such as CRH, which could act on the local BNST microcircuitry and through projections to downstream regions such as PVN (6).

## Conclusion

Maladaptive changes of BNST function underlie pathological anxiety disorders in humans (3). Here we identified 1) a novel electrophysiological mechanism suggesting CVMS activates ovBNST and 2) dysfunctional BNST CRH signaling represents a novel chronic stress-associated molecular signature. Activation of ovBNST coupled with neurochemical alterations in BNST_ALG_ may result in disturbances that underlie stress-related mood disorders. Electrophysiological and behavioral effects of CVMS were reversed by ovBNST PKA inhibition. Furthermore, optogenetic activation of ovBNST neurons mimicked the effects of CVMS by increasing negative valence behaviors associated with anxiety. Taken together, these data suggest that ovBNST is a critical component of the neural circuitry that underlies stress-induced mood disorders.

## Supporting information

Supplemental Figure 4 Part 1

Supplemental Figure 4 Part 2

Supplemental Figure 7 Part 1

Supplemental Figure 7 Part 2

Supplemental Figure 7 Part 3

## Acknowledgements

The authors would like to acknowledge Prof. Tracey J. Shors; Dr. Mimi Phan; Christine Yohn; Frederric Kelada; Kaci Shu; Bren Wu; Ashley Huang; Nicole Jallali; Andrew Dieterich; Ali Yasrebi; and Hannah Wang for helpful discussions and/or technical assistance.

