## Supplemental File for "Corticotropin-Releasing Hormone Signaling in the Oval Bed Nucleus of the Stria Terminalis Mediates Chronic Stress-Induced Negative Valence Behaviors Associated with Anxiety"

**Supplementary File**

**SUPPLEMENTARY MATERIALS AND METHODS:**

**Anxiety/Depression-like behavior tests:**

All behavioral tests were performed between 8:00 and 12:00 am. Mice were first put into the behavior room for at least 20 min adaptation. In total, 20 mice (n=10 for each group) were tested.

**- Sucrose Preference Test**

Sucrose preference test was performed starting the same day after completion of CVMS procedures to evaluate anhedonia, a core symptom of depression . Animals were first trained to drink 1% (w/v) sucrose solution for a 24 h adaptation period at Day 0. At 8:30 am in the morning of the testing day (Day 1), they were given free access to two bottles (each containing normal water and sucrose solution respectively). To avoid left/right preference, the order of bottle (left-right placement of water vs. sucrose) was alternated for each mouse during middle timepoint of both the adaptation and testing period. Bottles were weighed at the beginning and end of the testing 24 h test period. The percentage of sucrose solution relative to the total liquid consumed during this 24 h was used as an index of anhedonia.

**- Elevated Plus Maze (EPM)**

The EPM test evaluates anxiety-like behavior . Mice were placed in the central arena of a black plus-shaped maze at Day 2 after CVMS procedures, facing an open arm and were left to explore for 10 min. The duration and frequency of which open arms were explored was analyzed by video camera and processed by EthoVision (Noldus, Wageningen, The Netherlands).

**- Open-field Test**

The open-field test analyzes spontaneous exploratory activity and curiosity to a novel environment and is often used to evaluate anxiety-like behavior together with the EPM test . The **o**pen field apparatus consisted of a black floor area (40.5 cm × 40.5 cm), with a 37.5 cm high transparent wall. Mice were placed at Day 3 after CVMS procedures, in the center of the apparatus (center square) and monitored for 30 min. Data was collected and processed with EthoVision (Noldus, Wageningen, The Netherlands) and distance, duration of time that mice spent in the center, and the frequency of entry of the mice into the center area was documented and analyzed.

**- Novelty Suppressed Feeding Test**

The NSF test elicits competing motivations between the drive to eat and the fear of venturing into center of brightly lit arena), and thereby evaluates anxiety .

Mice were first deprived of food for 18 h (starting at 2:00pm Day 3 afternoon) and then placed into holding cages at 8:00 am on Day 4. After 60 min, the mice were placed into a novel, brightly lit (1200 lux) arena (16” × 20”) with a pellet of chow placed in the center of the arena affixed to a circular platform of white filter paper (10 cm). The time taken for the mice took to bite the food pellet for the first time was recorded as the latency to eat, at which point the pellet was immediately removed from the arena. Those mice that did not eat were assigned a latency of 360 s.

**- Forced Swimming Test**

The forced swimming test was performed as described before . Mice were placed at Day 5 after CVMS procedures, into a clear plastic container of 10 cm in diameter and 30 cm deep, filled two-thirds of the way with 23-26 °C water and were videotaped for the entire session. Mice were placed in the forced swim containers for 6 min, but only the last 4 min were scored, and processed with EthoVision (Noldus, Wageningen, The Netherlands).

**Basal Plasma Corticosterone Concentration Assay:**

After completion of the CVMS procedures, another 18 mice (n=9 from the Control and CVMS group) were quickly decapitated after anesthetization with euthasol (pentobarbital sodium; 150 mg/kg i.p.). Trunk blood samples were collected into heparin-coated tubes between 9-11:00 am, and then immediately chilled on ice and centrifuged at 4,000 rpm for 15 min at 4°C. Plasma was then stored at -80 °C for measuring plasma CORT using an enzyme-linked immunoassay kit (K014-H1; DetectX, Arbor Assays, MI).

**Real-time quantitative RT-qPCR analysis:**

After decapitation, the BNST tissue was dissected according to literature (AP +0.10 mm to -0.46 mm). Then, antero-dorsolateral BNST (BNSTadl) was punched out from 160 μm thickness sections and stored at -80°C. Total RNA and protein from BNSTadl was extracted using the RNA/Protein purification plus kit (48200; Norgen Biotek CORP, ON, Canada) and stored at -80°C.

mRNA expression was analyzed with eukaryotic 18S rRNA endogenous control (Taqman VIC^TM^ probe; Invitrogen, CA) as control primer for PCR amplification. cDNA was synthesized using SuperScript™ IV VILO™ Master Mix (with ezDNase™ enzyme) (11766050; Invitrogen, CA) as the reverse transcriptase. Q-PCR was performed with Taqman Fast Advanced Master Mix (4444557; ThermoFisher Scientific, CA), control probe and each target gene FAM Taqman Probe. The relative amount of target gene was calculated using the 2^-ΔΔCt^ method .

**Western blot study:**

Protein samples from the anterior-dorsolateral BNST (BNSTadl) of Control vs. CVMS stress mice were probed with an anti-phospho-PKA C (Thr197) primary antibody (rabbit, poly; 1:1000; 4781; Cell Signaling, Danvers, MA) and anti-PKA (C-α) primary antibody (rabbit, poly; 1:1000; 4782; Cell Signaling, Danvers, MA), as well as anti-STEP primary antibody (mouse, mono; 1:1000; 23E5; Cell Signaling, Danvers, MA), and then horseradish peroxidase (HRP)-conjugated goat-anti-rabbit and goat-anti-mouse IgG secondary antibodies (1:10,000; 31460, Thermo Fisher Scientific, Waltham, MA) respectively, operated on an iBind Flex Western device (Thermo Fisher Scientific, Waltham, MA). GAPDH was used as internal control (rabbit; 1:10,000; G9545; primary antibody purchased from Sigma-Aldrich, Saint Louis, MO). Immunoblot signals were then detected with an ECL chemiluminescence system (SuperSignal West Pico chemiluminescence Substrate, Pierce, IL) and imaged with myECL imager (Thermo Fisher Scientific, Waltham, MA). Immunoblots were quantified and density was calculated using ImageJ software (Wayne Rasband, NIMH, Bethesda, MD). Results were expressed as percentage of GAPDH.

**Brain tissue, immunohistochemistry and image acquisition:**

After behavioral tests were completed, 12 mice (n=6 for each CVMS or Control group) were anesthetized with isoflurane in individual cages and then perfused transcardially at Day 6 after CVMS procedures, with saline followed by 4% paraformaldehyde in PBS. Brains were cryoprotected in 30% sucrose before 40 μm thick sections were cut on a vibratome.

Immunohistochemical staining was performed according to standard procedures , with primary antibodies anti-CRH (rabbit, ab8901, 1:400; Abcam, MA), anti-c-fos (rabbit, 9F6, 1:800; Cell Signaling, MA), anti-PACAP (rabbit, ab216627, 1:700; Abcam, MA) and anti-STEP (mouse, 23E5, 1:500; Novus Biologicals, CO). For PACAP and STEP staining, mounted sections were first processed by antigen-retrieval through being heated in a citriate buffer (pH 6.0; Sigma, MO) in a microwave oven for 2 minutes followed by cooling down at least 30 min. Amplification of the signal was performed with biotinylated goat-anti-rabbit (A27035, 1:10000; Invitrogen, CA) or biotinylated goat-anti-mouse (A28176, 1:10000; Invitrogen, CA) IgG superclonal secondary antibodies, followed by avidin-biotin complex (1:50; PK6100, Vector Laboratories, CA). Chromogen development was performed with diaminobenzidine (1:50; SK-4100, Vector Laboratories, CA; 0.01% H2O2). Photographs were taken using Invitrogen EVOS FL Auto 1 Cell Imaging System (Invitrogen, CA). C-fos-, CRH-, PACAP- and STEP- immunopositive cells expressed in both the oval nucleus (ovBNST) and in the surrounding anterolateral dorsal region of BNST (BNSTad) (see the anatomy example shown in Fig. S4E) of the anterior BNST region (ranging from AP +0.10 mm to -0.46 mm) were counted bilaterally at a 20× magnification by an investigator blind to the experimental conditions.

**Immunofluorescence Study:**

To assess whether CRH cells co-localize with PACAP or STEP in the ovBNST, additional double immunofluorescence stainings were performed. The following antibodies and conditions were used: CRH (rabbit, ab8901, 1:400; Abcam, MA), PACAP (mouse, sc-166180, 1:200; Santa Cruz, TX) and STEP (mouse, sc-23892, 1:400; Santa Cruz, TX). After permeabilized with 0.1% Triton for 5 min, mounted BNST sections were washed with TBS for 5 min, then blocked in 10% normal goat serum and 0.3% Triton for 20 min, and then incubated in primary antibodies diluted in 2% normal goat serum and 0.3% Triton at room temperature for 1 h, then incubated overnight at 4 °C. The next day sections were washed and incubated for 2 h in Alexa Fluor Plus 555 (goat-anti-rabbit, A32732, 1:1000; Invitrogen, CA), or FITC (goat-anti-mouse, A16079, 1:1000; Invitrogen, CA). After brief rinse, sections were embedded in the ProLong diamond antifade mountant (P36961; Invitrogen, CA). Fluorescent signals were detected and photographs were collected using Invitrogen EVOS FL Auto 2 Cell Imaging System (Invitrogen, CA).

**Tissue preparation for brain slices patch clamp recording:**

Slices were prepared as described previously . Mice were quickly decapitated between 10:00–11:00 A.M. The brain was rapidly removed from the skull, and a block containing the BNST was immediately dissected and submerged in cold (4°C) oxygenated (95% O_2_, 5% CO_2_) high-sucrose artificial cerebrospinal fluid (aCSF) (in mM): 208 sucrose, 2 KCl, 26 NaHCO_3_, 10 glucose, 1.25 NaH_2_PO_4_, 2 MgSO_4_, 1 MgCl_2_, 10 HEPES, pH 7.3, 300 mOsm. Coronal slices (250 μm) were cut on a vibratome at 4°C. The slices were then transferred to an auxiliary chamber in which they were kept at room temperature (25°C) (recovery for 1-2 h) in aCSF consisting of the following (in mM): 124 NaCl, 5 KCl, 2.6 NaH_2_PO_4_, 2 MgCl_2_, 2 CaCl_2_, 26 NaHCO_3_, 10 glucose, pH 7.3, 310 mOsm until recording. A single slice was transferred to the recording chamber mounted on an Olympus BX51W1 upright microscope. The slice was then continually perfused with warm (35°C), oxygenated aCSF at 1.5 ml/min. Targeted neurons were viewed with an Olympus 40x water-immersion lens.

**Electrophysiological Recordings (M-current and mEPSC):**

Electrophysiology was performed as described previously . In BNST slices, standard whole-cell voltage-clamp patch recording procedures and pharmacological testing were performed. Recordings were performed using pipettes made of borosilicate glass and pulled using a PC-10 Puller (Narishige, Japan). Axopatch 700B amplifier, Digidata 1322A Data Acquisition System, and pCLAMP software (version 10.2; Molecular Devices, Sunnyvale, CA) were used for data acquisition and analysis.

All recordings were restricted to the ovBNST (anatomy shown in Fig 2A; with bregma -0.26 mm in Swanson Brain Atlas, interaural 8.74 mm) . Recordings of ovBNST neurons were based on both 1): the anatomical criteria: located within the anterodorsolateral BNST (located dorsal to the halfway point between the tip of lateral ventricle and top of anterior comissure); and (2): morphological criteria: medium-sized somata. Only neurons that qualified for the criteria were recorded and analyzed subsequently.

Current-voltage (*I-V*) plots were generated by voltage steps from –50 to –140 mV at 10 mV increments applied at 1-s intervals from a holding potential of –60 mV. From this protocol, the input resistance was determined from the slope of the *I-V* plot in the range between –60 and –80 mV. Input resistance, series resistance, and membrane capacitance were all monitored throughout the experiments. Only cells with stable series resistance (< 30 MΩ; < 20% change over the course of the recording) and suitable input resistance (> 500 MΩ) were used for analysis.

To record M-currents, pipettes (3-5 MΩ resistance) were filled with internal solution containing (in mM): 10 NaCl, 128 K-gluconate, 1 MgCl_2_, 10 HEPES, 1 ATP, 1.1 EGTA, 0.25 GTP; pH 7.3, 300 mOsm. 1 M TTX was included in the recording ACSF to block Na^+^-spike–dependent synaptic inputs. All the drugs used here were purchased from Tocris (Minneapolis, MN). Under the voltage clamp, a standard deactivation protocol (Fig. 2C) was used to measure K^+^ currents elicited during 500-ms voltage steps from –30 to –75 mV in 5-mV increments after a 300-ms prepulse to –20 mV. The amplitude of M-current relaxation or deactivation was measured as the difference between the initial (< 10 ms) and sustained current (> 475 ms) of the current trace under control condition (TTX only; 1 μM; 5 min). After baseline recording (~ 5 min), deactivation protocol was repeated twice and averaged for analysis.

To study excitatory synaptic transmission in the ovBNST, pharmacologically isolated mEPSCs were recorded as described . Picrotoxin (PTX, 50 μM) was added to block inhibitory synaptic transmission mediated by GABA_A_ receptor, and meanwhile D-APV (50 μM) was added to block currents mediated by NMDA receptors. In addition, TTX (1 μM) was included to block action potential. Internal solution contained (in mM): 40 CsCl, 10 HEPES, 0.05 EGTA, 1.8 NaCl, 3.5 KCl, 1.7 MgCl_2_, 2 Mg-ATP, 0.4 Na_4_-GTP, 10 phosphocreatine, and 5 N-(2,6-Dimethylphenylcarbamoylmethy)triethylammonium, and was adjusted to pH 7.3, 280-290 mOsm. All the drugs used here were also purchased from Tocris (Minneapolis, MN). After a stable baseline recording of approximately 5-min period, mEPSC was continuously recorded for around 10 min. The mEPSC properties recorded during the last 5 min period were compared.

To test the possibility of PKA-mediated effects of CVMS on the electrophysiological properties, several coronal slices of BNST from chronic stress mice were incubated with 10 μM H89 (the selective PKA inhibitor; from Tocris, Minneapolis, MN) in the ACSF for at least 30 min ahead at room temperature before transferred to the recording chamber for current recording.

**Comparison of electrophysiological recordings (M-current and mEPSC) and immunohistochemistry in posterior BNST (pBNST):**

Tissue preparation and all the slice recording processes and methods for brain slice patch clamp recordings in the pBNST were as same as in the main method. Posterior BNST region were included ranging from bregma AP – 0.51mm to – 1.08 mm . Coronal slices (250 μm) were cut. We applied the following criteria for patch clamping recordings: 1) only cells within the above anatomy location criteria of pBNST will be considered; 2) within pBNST, we will select the principle nucleus (BNSTpr) region, which ranges from bregma AP -0.7 to -0.9 mm (refer to Fig. 1G in ; 3) within the BNSTpr region, we select only the medium-sized cells.

All the immunohistochemistry methods used and details are the same as in the main method.

**Local cannula drug infusion of PKA-selective antagonist H89 into BNST:**

To test the possibility of PKA-mediated behavioral effect of CVMS, either CVMS mice (n=7-8) or Control mice (n=7-8) were anaesthetized with 1.5-3.0% isoflurane and placed on a stereotaxic apparatus (Kopf Instruments, Tujunga, CA) with a heating pad (T/Pump, Stryker, Kalamazoo, MI) underneath. They were bilaterally implanted with a guide cannula (C315G/SPC, Plastics1, Roanoke, VA) directly into ovBNST (bregma AP +0.2 mm, ML 1.0 mm, DV -4.1 mm) (example picture shown in Fig. 6B). Using dental cement, a dummy cannula (C315DC/SPC, Plastics1, Roanoke, VA) was capped on it to help protect guide cannula and prevent debris from entering. Mice were put into heated home cage to allow recovery from anesthesia for at least 20-30 min before put back into colony room to allow further recovery for at least 1 week before drug infusion starts.

During drug infusion, the dummy cannula was gently taken out first, then an internal cannula (also called infusion cannula; C315I/SPC, Plastics1, Roanoke, VA) was slowly inserted directly into guide cannula. Then a Hamilton syringe (HSYR-1 Syringe 86200, Plastics1, Roanoke, VA) connected with a segment of PE tubing (C313CT/PKG, Plastics1, Roanoke, VA) was used to gently infuse H89 (Tocris, Minneapolis, MN; 25 nM/dissolved in 0.5 μl saline) at a rate of 0.05 μl/min into ovBNST (bregma AP +0.2 mm, ML 1.0 mm, DV -4.1 mm) , with the help of a 0.58 mm diameter flexible connector assembly (C313C/SPC, Plastics1, Roanoke, VA) connected to the internal cannula. Since this connector secures the internal cannula onto guide cannula with the help of captive collar, this permits the whole infusion process to proceed smoothly while the mice is undisturbed and able to freely move in the cage, hence preventing any unnecessary stress if mice would have been held or restrained otherwise. After the completion of H89 infusion, the internal cannula was kept in place for at least 5-10 min before gently taken out. Then the dummy cannula was put back and mice was gently placed back into its home cage. H89 was chronically infused for a continuous 7 days to allow fully penetration into the BNST tissue. Daily CVMS exposure was continuously present during the 7-day H89 chronic drug infusion period.

**AAV-virus stereotactic injection and optic-fiber implantation:**

Adeno-associated virus (AAV) vectors were serotyped with AAV5 coat proteins and packaged by University of North Carolina Vector Core (Chapel Hill, NC) and Add Gene (Watertown, MA). All surgical procedures were performed aseptically. Male Drd1a-Cre transgenic mice (that express Cre-recombinase from a Drd1a promoter) (8-10 week age) were anaesthetized with 1.5-3.0% isoflurane and placed on a stereotaxic apparatus (Kopf Instruments, Tujunga, CA) with a heating pad (T/Pump, Stryker, Kalamazoo, MI) underneath. They were bilaterally injected with each side at least 0.35 μl of Cre-dependent either AAV5-EF1α-DIO-ChR2 (H134R)-eYFP virus (ChR2:ovBNST; as ChR2) or control AAV5-EF1α-DIO-eYFP-virus (eYFP:ovBNST; as Control) (from University of North Carolina at Chapel Hill, NC; titer 2 X10^12 particles/ml) into ovBNST (AP +0.2 mm, ML 1.0 mm, DV -4.1 mm; relative to bregma) of Drd1a-Cre mice (to ensure the virus to be selectively expressed in the ovBNST), using a Nanoject III Programmable Nanoliter Injector (Drummond Scientific Company, Broomall, PA). The injection rate was set to 2 nl/s by the Nanojector program.

After completion of virus injection each side, the injector needle remained in place for at least additional 5-7 min before slowly withdrawal to avoid upward flow and allow stable penetration of virus into brain tissue. Then a pre-polished fiberoptic cable (FT200UMT; 200μm Core; Thorlabs, Newton, NJ) connected through ceramic ferrule (CFLC230-10; diameter 1.25 mm, length 6.4 mm, Thorlabs; Bore size 230 μm) was slowly implanted directly above ovBNST into each side. Adhesive glue was first applied surrounding the ferrule and then dental cement was added to secure the ferrule to the skull. Finally the incision was closed using tissue adhesive. Mice were put into a warm cage to facilitate recovery from anesthesia for at least 30 min before they were placed back into the colony room. They were allowed to recover for at least additional 3 weeks before subsequent behavioral experiments conducted to allow enough time for opsin virus expression.

**Light Delivery and optogenetic-controlled behavioral tests:**

For optogenetic ChR2 stimulation, 3-5 mW of blue light (95-159 mW/mm­^2^) was generated by a 473 nm DPSS laser system (MBL-III 473; Opto Engine LLC, Midvale, UT) and bilateral delivered through a fiberoptic patch cord made from pre-polished firberoptic cable (FT200UMT; 200 μm Core; Thorlabs, Newton, NJ) connected through a ceramic ferrule (CFLC230-10; diameter 1.25 mm, length 6.4 mm, Thorlabs; Bore size 230 μm) and a stainless steel ferrule multimode connector (F12774, 230 μm; Fiber Instrument Sales, Inc., Oriskany, NY). Blue laser output was controlled using a pulse generator (Master-8; A.M.P.I., Jerusalem, Israel) to deliver 5-ms light pulse trains at 10Hz frequency.

For the optogenetic-controlled EPM test, each of the light OFF-ON-OFF session consists of a 5-min duration; and for the optogenetic-controlled Open Field test, each of the light OFF-ON-OFF session consists of a 10-min duration. Light OFF or ON button was precisely controlled manually by the investigator through the Master 8 pulse generator (A.M.P.I., Jerusalem, Israel).

**SUPPLEMENTARY RESULTS:**

1. **Basal plasma corticosterone (CORT) concentration:**

Basal CORT concentration between CVMS group and the Control group was compared and shown in Fig. S1. Significantly higher basal CORT concentration was found in the CVMS group (2059 ± 316.7 pg/ml; n=10) vs. Control group (924.5 ± 123.5 pg/ml; n=10; p<0.01).

2. **Anxiety/Depressive-like behavior induced by CVMS:**

6 weeks of CVMS resulted in a significant percent decrease in body weight (CVMS: -12.60±0.58%; n=9) compared with non-stress controls (Control: 18.62±4.14%; n=6; ANOVA F(1,13)=84.908, p<0.001). As shown in Fig.S1, significantly higher basal CORT levels in the CVMS group (2059±316.7 pg/ml; n=10) was found relative to the control group (924.5±123.5 pg/ml; n=10; ANOVA F(1,18)=12.281, p<0.01).

In the Elevated Plus Maze (EPM), we found a significantly decreased open arm duration (314.9±21.44 s vs. 415.7±27.74 s; ANOVA F(1,18)=8.262; p=0.01) in the CVMS group (n=10) compared with controls (n=10) (Fig.1B). Entry frequency into open arms was not different (86.80±7.36, n=10 vs. 73.3±5.84, n=10 in the control group; ANOVA F(1,18)=2.065; p=0.17) (Fig.1C). In the sucrose preference test (SPT), preference was significantly decreased in the CVMS group (47.99±6.01%; n=10) compared with the control group (78.61±2.11%; n=10; ANOVA F(1,18)=23.118; p<0.001) (Fig.1D). Third, in open field (OF) test, CVMS (n=10) resulted in a significant decrease in center distance (605.3±38.39 cm vs. 1878±228.6 cm; ANOVA F(1,18)=30.155; p<0.001) (Fig.1E), center duration (61.85±5.05 vs. 181.1±14.65 s; ANOVA F(1,18)=59.24; p<0.001) (Fig.1F) and center entry frequency (52.10±3.78 vs. 100.8±10.36; ANOVA F(1,18)=19.50; p=0.001) (Fig.1G) compared with controls (n=10). Fourth, in the Novelty Suppressed Feeding (NSF) test, CVMS (n=10) significantly increased the latency to eat (137.8±17.7 s vs. 68.9±11.86 s; ANOVA F(1,18)=10.460; p<0.01) (Fig.1H). Finally, CVMS had no significant effect on immobility in the Forced Swim Test (FST) (359.6±0.11 s, n=9 vs. 316.9±22.10 s, n=11 in Control group; ANOVA F(1,18)=3.035; p=0.08) (Fig.1I).

3. **Ephys property changes induced by CVMS:**

CVMS significantly depolarized the resting membrane potential (RMP) (-61.04±2.03 mV [n=5] compared to -68.00±0.79 mV in Control [n=6]; t-test t=3.192, p=0.023) (Fig.2F) and increased the input resistance (Rin) (1.95±0.08 GΩ [n=5] vs. 1.18±0.20 GΩ in Control [n=5]; t-test t=3.632, p=0.014) (Fig.2G) in ovBNST neurons.

CVMS significantly increased the average amplitude of mEPSCs (Fig.2H-J) from 13.88±0.81 pA (Control, n = 6) to 20.22±2.00 pA (CVMS, n = 5; t-test t=3.141, p = 0.012). There was no change in the mEPSC frequency (Control: 6.31±1.36 HZ [n=6] vs. CVMS: 4.97±1.02 HZ [n = 5]; t-test t=0.790, p=0.451 (Fig.2J).

**4. IHC study:**

The number of c-fos-immunoreactive cells number in BNSTadl and ovBNST were significantly increased in CVMS mice relative to control mice (BNSTadl: 2025±98 [n=6] in CVMS vs. 1097±158 in control [n=6]; ANOVA F(1,10)=24.896, p<0.01; ovBNST: 1130±54 in CVMS [n=6] vs. 490±58 in control [n=6]; ANOVA F(1,10)=65.29, p<0.001) (Fig.3.2A-B). Similar to cFos, CVMS increased the number of CRH+ cells in BNSTadl and ovBNST relative to control (BNSTadl: 3277±90 [n=6] vs. 2315±103 [n=6] in control; ANOVA F(1,10)=49.78, p<0.01; ovBNST: 1883±133 [n=6] vs. 1100±59 [n=6] in control; ANOVA F(1,10)=28.93; p<0.01) (Fig.3.2C-D).

The cell number of PACAP in the BNSTadl was 4113±99 in CVMS [n=6] vs. 2790±113 in control [n=6]; (ANOVA F(1,10)=77.89, p<0.01); and the cell number of PACAP in the ovBNST was 2390±103 in CVMS group [n=6] vs. 1318±33 in control [n=6]; (ANOVA F(1,10)=98.36, p<0.001) (shown in Fig.3.2.E and F)). The cell number of STEP in the BNSTadl was 2164±135 in CVMS [n=5] vs. 3228±229 in control group [n=5]; (ANOVA F(1,8)=16.063, p<0.01); and the cell number of STEP ovBNST: 1228±121 in CVMS [n=5] vs. 2134±162 in control group [n=5]; (ANOVA F(1,8)=20.099, p<0.01) (Fig.3.2G and H).

**5. qPCR study:**

CVMS significantly increases CRH mRNA in BNSTadl (131.2±5.1%; n=7) compared with control (102.9±9.1%; n=9; ANOVA F(1,14)=6.303, p=0.018) (shown in Fig 4B). Similarly, PACAP expression in BNSTadl of CVMS mice was also significantly increased (129.7±7.1% vs. 102.4±9.1 in control; both n=7; ANOVA F(1,12)=5.597, p=0.037) (Fig.4C). By contrast, STEP expression in BNSTadl of CVMS mice was significantly decreased (71.3±7.0% vs. 105.7±12.3% in control; both n=9; ANOVA F(1,16)=5.877, p=0.031) (Fig.4D). Interestingly, we also found that CRHR1 expression was significantly increased in CVMS mice (128.3±8.2%; n=8 vs. 102.6±8.1%; n=9 in control; ANOVA F(1,15)=4.985, p=0.041) (Fig.4E), whereas expression of CRHR2 was unchanged (133.3±27.7%; n=9 in CVMS vs. 97.0±31.6%; n=7 in control; ANOVA F(1,14)=0.750, p=0.403) (Fig.4F).

**6. Western blot study:**

While no significant differences were found in expression levels of total PKA in BNSTadl of CVMS mice (80.91±2.52% of GAPDH; n=7) vs. control (85.75±2.16% of GAPDH; n=7; *t* test t=1.46, p=0.171) (Fig.4I), p-PKA expression in BNSTadl is increased by CVMS (71.19±10.19% of GAPDH, n=7 vs. 14.48±4.41% of GAPDH, n=7 in control; *t* test t=5.105, p=0.0003) (Fig.4J), indicating increased activation of PKA in BNSTadl of CVMS mice.

There is no significant difference in expression of the membrane isoform STEP61 in BNSTadl of CVMS mice (18.92±7.12% of GAPDH; n=7) vs. control (29.89±7.52% of GAPDH; n=7; *t* test t=1.060, p=0.31) (Fig.4L), expression of the cytosolic isoform STEP46 in BNSTadl is decreased from 87.55±15.84% of GAPDH in control mice (n=7) to 49.31±5.27% in CVMS mice (n=7; *t* test t=2.292, p=0.026) (Fig.4K), suggesting that only cytosolic STEP46 isoform expression is decreased in the BNST of CVMS mice.

**7. Effects of PKA-selective antagonist H89 infused into ovBNST of CVMS mice on ephys properties and anxiety-like behavior:**

At –40 mV, the outward M-current peak value was increased from 46.28±16.93 pA (n = 6) in CVMS slices to 100.76±16.68 pA (n=6; *t*-test t=2.292, p=0.045) in CVMS slices pre-incubated with H89 (Fig.5A). Increased amplitude of mEPSCs in ovBNST of CVMS mice (20.22±2.00 pA; n=5) was significantly decreased to 13.36±1.24 pA (n = 4; *t*-test t=2.913, p=0.025) when slices were preincubated with the selective PKA antagonist H89 for 30 min (Fig.5B). By contrast, mEPSC frequency in ovBNST was not different between CVMS mice (4.97±1.02 Hz; n=5) and CVMS+H89 mice (4.45±2.26 Hz; n=4; *t*-test t=0.210; p=0.843) (Fig.5C).

In EPM, BNST H89 infusions significantly increased the duration of CVMS mice in open arms (CVMS+H89: 457.8±38.4 s, n=6; vs. CVMS: 314.9±21.4 s, n=10; ANOVA F(1,14)=12.528; p=0.011) (Fig.6C). There were no significant differences in frequency of open arm entries between the two groups: Stress+H89: 104.7±17.3 (n=6), vs. Stress: 86.8 ±7.4 (n=10; ANOVA F(1,14)=1.212; p=0.374) (Fig.6D). In OF, H89 significantly increased the distance that CVMS mice traveled in the center (CVMS+H89: 1890±289 cm, n=7; vs. CVMS: 605±38 cm, n=10; ANOVA F(1,15)=27.962; p<0.01) (Fig.6E), the duration of time that CVMS mice spent in the center (CVMS+H89: 260.7±42.1 s, n=7; vs. CVMS: 61.9±5.1 s, n=10; ANOVA F(1,15)=31.752; p<0.01) (Fig.6F), and the frequency of center entries (CVMS+H89: 90.6±9.5, n=7; vs. CVMS: 52.1±3.8, n=10; ANOVA F(1,15)=18.121; p<0.01) (Fig.6G). In the SPT, H89 significantly increased sucrose preference in CVMS mice (CVMS+H89: 65.9±6.2%, n=6 vs. CVMS: 46.7±5.1%, n=10; ANOVA F(1,19)=4.43; p=0.035) (Fig.6H). Finally**,** in NSF, H89 significantly decreased the latency of CVMS mice to eat (CVMS+H89: 61.8±23.2 s (n=5); vs. CVMS: 137.8±17.7 s (n=10); ANOVA F(1,19)=4.434; p=0.025) (Fig.6I).

**8. Comparison of mRNA expression for KCNQ2, KCNQ3 and KCNQ5 subunit:** mRNA expression for KCNQ2, KCNQ3 and KCNQ5 subunit (that encodeKv7M-channel composition) in the anterior-dorsolateral BNST (BNSTadl) from both Control group and CVMS group was also analyzed and compared. Interestingly, as shown in Fig. S2, no significant difference was found for all these 3 genes that we examined (all p>0.05, CVMS vs. Control).

9. **Comparison of immunoreactivity of c-fos, CRH, PACAP and STEP in the**

**anterolateral dorsal region of BNST (adBNST):**

Lower magnification of comparison of immunostaining patterns of CRH (Fig. S3A and B), c-fos (Fig. S3C and D), PACAP (Fig. S3E and F) and STEP (Fig. S3G and H) in the anterior-dorsolateral BNST (BNSTadl) from Control mice vs. CVMS mice was shown in Fig. S3. Higher magnification of immunostaining pattern of c-fos (Fig. S4.1A and B), CRH (Fig. S4.1C and D), PACAP (Fig. S4.1E and F) and STEP (Fig. S4.1G and H) in the BNSTadl was compared and shown in Fig. S4.1, with black dots highlighting the anatomy boundary within which oval nucleus of BNST (ovBNST) is located in the BNSTadl.

We also quantified immunoreactivity (IR) of the above markers (c-fos, CRH, PACAP and STEP) expressed in the antero-dorsal region of BNST (adBNST), the region closely surrounding oval nucleus within the BNSTadl (example shown in Fig. S4.2E):

1) CRH-IR cells in the adBNST region did not differ significantly (1393 ± 92) in CVMS group, compared to those (1215 ± 53) in Control group (n=6; ANOVA F(1,10)=2.792; p>0.05) (shown in Fig. S4.2A);

2) Similarly, c-fos-IR cell number in the adBNST region did not differ significantly in CVMS group (895 ± 55), compared to those in Control group ((607 ± 111; n=6; ANOVA F(1,10)=5.428; p>0.05) (Fig. S4.2B);

3) Again, PACAP-IR cells in the adBNST region did not differ significantly (1393 ± 92) in CVMS group, compared to those (1215 ± 53) in Control group (n=6; ANOVA F(1,10)=3.287; p>0.05) (shown in Fig. S4.2C);

4) Likewise, as shown in Fig. S4.2D, STEP-IR cells in the adBNST region did not differ significantly (936 ± 46) in CVMS group, compared to those (1094 ± 86) in Control group (n=5; ANOVA F(1,8)=2.644; p>0.05).

Together, for all of the 4 markers that we examined, there is no significant change in the cell number expressed in the adBNST compared between Control vs. CVMS group.

**10. Immunofluorescence co-localization of CRH with PACAP and STEP in the ovBNST:**

Since previous study has shown STEP and CRH immunoreactivity co-localize in the neurons of the oval nucleus of BNST (ovBNST) , we next sought to confirm the expression pattern of these two neuropeptides in the ovBNST by immunofluorescence study. Consistently, we also found almost complete co-localization of CRH with STEP in the ovBNST (shown in Fig. S5A-C, CRH in red and STEP in green color).

Since high density of PACAP expression was found in the ovBNST , and since PACAP and CRH peptide signaling are suggested to be integrated to regulate BNST activity, we also sought to explore the expression pattern of these two neuropeptides in the ovBNST. Similarly, a high percentage of co-localization of CRH with PACAP neuropeptides was also found in the ovBNST (shown in Fig. S5D-F, CRH in red and PACAP in green color).

Together, our results show that both STEP and PACAP shows a pattern of highly co-localization with CRH in the ovBNST.

**11. Comparison of electrophysiological properties and immunohistochemistry in the pBNST:**

Since literature has suggested the posterior medial region of the BNST (BNSTpm) is an area that is involved in limiting HPA responses to acute stress , we also set out to examine changes in the electrophysiological properties and immunohistochemistry in the pmBNST after CVMS. Posterior BNST region were included ranging from AP – 0.51mm to – 1.08 mm .

1) **CVMS does not affect M-current or mEPSC in the pBNST:**

Recordings of posterior BNST neurons focus on the medial area of the posterior BNST (BNSTpm) neurons, mainly on the principal nucleus (BNSTpr) region, located lateral to the stria medularis (sm) (example anatomy was shown in Fig. S6A, same as previously reported in ), with coordinates ranging from bregma AP -0.7 to -0.9 mm (please refer to Fig. 1G in ).

Interestingly, CVMS has no significant effect on the M-current (F(1,10)=1.608 , p=0.234) recorded at any voltage tested. The peak current at -35 mV is 190.02 ± 52.62 pA in the Control group (n=7) vs. 231.90 ± 28.95 pA (n=5; p>0.05) in the CVMS group, respectively (Fig. S6B).

Consistently, there was no significant change in either the average amplitude (shown in Fig. S6C) (Control: 13.0 ± 0.38 pA, n= 4; vs. CVMS: 14.1 ± 0.56 pA, n=4; p=0.162) or frequency (shown in Fig. S6D) (Control: 5.13 ± 1.32 HZ; n=4; vs. CVMS: 4.45 ± 1.28 HZ; n = 4; p =0.727) of mEPSCs compared between these two groups.

**2) CVMS has no significant effects on the number of c-fos, CRH, PACAP and STEP-immunoreactive (IR) cells expressed in the pBNST:**

Comparison of immunostaining patterns of c-fos (A,C for Control vs. B,D for Stress), CRH (E,G for Control vs. F,H for Stress), PACAP (I,K for Control vs. J,L for Stress) and STEP (M,O for Control vs. N,P for Stress) in the pBNST was shown in Fig. S7.1.

As shown in Fig. S7.2A, c-fos-immunoreactive (IR) cell number in the pBNST was 1245 ± 164 in CVMS group vs. 1048 ± 120 in Control group (both n=6; p=0.360).

Similarly, CRH-IR cell number in the pBNST was 4335 ± 168 in CVMS group vs. 3978 ± 122 in Control group (both n=6; p=0.120) (shown in Fig. S7.2B).

Again, PACAP-IR cell number in the pBNST was 4693 ± 100 in the CVMS group vs. 4950 ± 242 in the Control group (both n=6; p=0.361) (shown in Fig. S7.2C).

Likewise, STEP-IR cell number in the pBNST was 5502 ± 141 in the CVMS group (n=5) vs. Control 5690 ± 312 (n=6; p=0.603) (shown in Fig. S7.2D).

Taken together, the above results indicated that for all the above 4 immuno-markers that we examined, there is no significant change in the cell number expressed in the pBNST compared between CVMS group vs. Control group.

**3) qPCR comparison of mRNA expression of CRH, PACAP and STEP, and CRH receptor CRHR1 and CRHR2 in the pBNST of control vs. CVMS mice:**

As shown in Fig. S7.3A, CVMS has no significant effect on the CRH mRNA expression in the pBNST (102.6 ± 8.7%; n=8) compared with Control group (106.2 ± 11.0%; n=9; p=0.800). Similarly, PACAP mRNA expression in the pBNST of CVMS group was not significantly changed (107.5 ± 14.6%) compared with that in the Control group (110.1 ± 16.6%; both n=9; p=0.909) (Fig. S7.3B). Again, CVMS has no significant effect on the STEP mRNA expression in the pBNST (109.6 ± 15.1%; n=8) vs. in the Control group (107.1 ± 15.0%; n=7; p=0.909) (Fig. S7.3C).

For the two CRH receptor subtypes that we examined, both mRNA expression of CRHR1 (107.2 ± 13.5% in CVMS (n=9) vs. 103.3 ± 8.3% in Control (n=9; p=0.809)) (Fig. S7.3D) and CRHR2 (112.8 ± 18.6% in CVMS (n=9) vs. 109.7 ± 15.7% in Control (n=9; p=0.901)) (Fig. S7.3E) has no significant change compared between the two groups.

Taken together, for all of the above 5 genes expressed in the pBNST (CRH, PACAP and STEP, and CRH receptor CRHR1 and CRHR2) that we examined, CVMS has no significant effect on their mRNA expression.

**12. Effects of PKA-selective antagonist H89 on the properties of M-current and mEPSC in the ovBNST of Control mice:**

BNST coronal slices from Control mice were also pre-incubated with 10 μM PKA-selective antagonist H89 (Control+H89) for at least 30 min to determine its effect on M-current and mEPSC properties. As shown in Fig. S8A, for M-current that we recorded in the ovBNST, no significant difference was found between Control group (n=6) vs. Control+H89 group (n=4) (group effect F(1,8)=0.021, p=0.888). M-current peak amplitude at -35 mV was: Control: 202.6 ± 39.9 pA (n=6) vs. Control+H89: 223.0 ± 33.0 pA (n=4; p=0.726).

Similarly, H89 has no significant effects on mEPSC properties recorded in the ovBNST. Amplitude of mEPSC (shown in Fig. S8B) was: Control 13.9 ± 0.8 pA (n=6) vs. Control+H89 14.7 ± 0.7 pA (n=4; p=0.517). Frequency of mEPSC (shown in Fig. S8C) was: Control 6.3 ± 1.4 Hz (n=6) vs. Control+H89 5.4 ± 0.5 Hz (n=4; p=0.534).

Together, the above results indicate that H89 has no significant effect on the properties of M-current and mEPSC in the ovBNST of Control mice.

**13. Effects of PKA-selective antagonist H89 infused into ovBNST on the anxiety/depressive-like behavior of Control mice:**

**1)** In the sucrose preference test, H89 has no significant effect on the sucrose preference intake percentage of Control mice either: Control+H89: (70.4 ± 7.5%, n=7); vs. Control: (75.1 ± 3.0%, n=15) (p=0.575) (shown in Fig. S9A).

**2)** As shown in Fig. S9B, in the EPM test, H89 has no significant effect on the duration time that the Control mice spent in the open arm (Control+H89: 430.6 ± 88.1 s, n=7; vs. Control: 423.7 ± 28.7 s, n=12; p=0.942) in the EPM test; similarly, there is no significant difference for the frequency that mice entered into the open arm compared between the two groups: Control+H89: 79.9 ± 20.3 (n=7), vs. Control: 70.6 ± 5.96 (n=11; p=0.673) (shown in Fig. S9C).

**3)** In the open field test, similarly, H89 does not significantly affect the distance that the Control mice traveled in the center of Open Field (Control+H89: 1957 ± 201.0 cm, n=7; vs. Control: 1870 ± 223.4 cm, n=13; p=0.775) (Fig. S9D); again, H89 has no significant effect on the duration that the Control mice spent in the center of Open Field: (Control+H89: 243.7 ± 47.6 s, n=6; vs. Control: 170.7 ± 16.8 s, n=13; p=0.196) (Fig. S9E); likewise, H89 does not significantly change the entry frequency that Control mice spent in the center of Open Field: (Control+H89: 87.1 ± 8.1, n=7; vs. Control: 98.2 ± 10.1, n=13; p=0.404) (Fig. S9F).

**4)** In the NSF test, likewise, H89 has no significant effect on the latency of Control mice to eat food pellet: latency time for Control+H89: 64.2 ± 38.4 s (n=5); vs. Control: 62.6 ± 8.7 s (n=15; p=0.952) (Fig. S9G).

Taken together, the above results suggest that PKA-selective antagonist H89 has no significant effect on the anxiety or depressive-like behavior of control mice when infused into ovBNST.

**14. Additional working model:**

**Since** our western blot result has indicated decreased expression of cytosolic STEP46 isoform after CVMS, whereas membrane isoform STEP61 has no change; given that PKA could phosphorylate and thus inactivate STEP activity , we then propose an additional working model in which under basal status, unphosphorylated (and active) STEP de-phosphorylates M-channel subunit and AMPA receptor (GluR1) subunit on the postsynaptic membrane, facilitating AMPAR (GluR1) internalization . After CVMS, activation of PKA will initiate STEP phosphorylation and its inactivation , which will in turn release a phosphorylation brake (shown in Fig. S10) and consequently result in extra-phosphorylation of both membrane M-channel and surface GluR1 (and more GluR1 surface trafficking and expression) . This will lead to a concomitantly decreased M-current and increased mEPSC amplitude . In this way, these two parallel pathways (direct phosphorylation by PKA and indirect release of phosphorylation brake through inactivated STEP) converge together to mediate hyper-phosphorylation of M-channel and AMPA receptor GluR1 subunit.

**Supplemental Figure Legends:**

**Fig. S1:** Basal plasma corticosterone (CORT) comparison between CVMS vs. Control group.

**Fig. S2:** Comparison of mRNA expression level of KCNQ channel subunit KCNQ2, KCNQ3 and KCNQ5 in the anterodorsal of BNST (BNSTadl) between CVMS vs. Control group.

**Fig. S3:** Lower magnification (4 times) comparison of the immunostaining pattern of c-fos, CRH, PACAP and STEP in the antero-dorsolateral BNST (BNSTadl).

**Fig. S4.1**: Comparison of immunostaining pattern of c-fos, CRH, PACAP and STEP in the antero-dorsolateral BNST (BNSTadl), with black dots highlighting the anatomy boundary within which oval nucleus of BNST (ovBNST) is located in the anterodorsal lateral region of the BNST (BNSTadl).

c-fos comparison: Immunostaining pattern of c-fos compared in the ovBNST of (A) Control (left) vs. (B) CVMS (right) mice;

CRH comparison: Immunostaining pattern of CRH compared in the ovBNST of (C) Control (left) vs. (D) CVMS (right) mice;

PACAP comparison: Immunostaining pattern of PACAP compared in the ovBNST of (E) Control (left) vs. (F) CVMS (right) mice;

STEP comparison: Immunostaining pattern of STEP compared in the ovBNST of (G) Control (left) vs. (H) CVMS (right) mice.

ic: internal capsule; ac: anterior commissure.

**Fig. S4.2:** Comparison of CRH (A), c-fos (B), PACAP (C) and STEP (D)-immunoreactive (IR) cell number in the anterodorsal region closely surrounding ovBNST, that is adBNST (Fig. S4E) from CVMS mice vs. Control mice. Fig. 4E shows anatomy example of the anterolateral dorsal region of BNST (adBNST), the region closely surrounding oval nucleus in the BNSTadl.

NS: non-significant different (p>0.05).

**Fig. S5:** Immunofluorescence co-staining (C and F) of CRH (red A and D) and PACAP (green B), STEP (green E) signaling in the oval nucleus of BNST (ovBNST). Scale bar: 50 μm.

**Fig. S6:** Anatomy example of principal nucleus (BNSTpr) region in the medial area of the posterior BNST (BNSTpm) (mainly located lateral to the stria medularis (sm)) is shown in Fig. S6A. Comparison of electrophysiological properties (including M-current (B) and mEPSC amplitude and frequency (C and D)) in the principle nucleus (BNSTpr) of posterior medial region of the BNST (BNSTpm) of Control vs. CVMS group.

**Fig. S7.1:** Comparison of c-fos (A,B,C and D), CRH (E,F,G and H), PACAP (I,J,K and L) and STEP (M,N,O and P)-immunostaining pattern in the posterior medial region of the BNST (BNSTpm) of Control vs. CVMS group.

**Fig. S7.2:** Comparison of c-fos (A), CRH (B), PACAP (C) and STEP (D)-immunoreactive (IR) cell number in the posterior medial region of the BNST (BNSTpm) of Control vs. CVMS group.

**Fig. S7.3**: qPCR comparison of CRH, PACAP, STEP, CRHR1, CRHR2 in the the posterior medial region of the BNST (BNSTpm) of Control vs. CVMS group.

**Fig. S8:** PKA-selective antagonist H89 was infused into the oval nucleus of BNST (ovBNST) of Control mice to examine its effect on the properties of M-current and mEPSC.

(A): I-V curve of outward M-current shows that PKA-selective antagonist H89 (n=6 cells) infused into the ovBNST of Control mice (Control+H89, n=4 cells) has no effect on the M-current of Control mice (n=6 cells) at all the membrane voltage examined (all p>0.05) ranging from -75 mV to -25 mV.

(B): H89 infused into the ovBNST of Control mice (Control+H89, n=4 cells) has no effect on the mEPSC amplitude of Control mice (n=6 cells) (p>0.05).

(C): H89 infused into the ovBNST of Control mice (Control+H89, n=4 cells) had no significant effect on the mEPSC frequency of Controlmice (n=6 cells) (p>0.05).

**Fig. S9:** **PKA-selective antagonist H89 was infused into the oval nucleus of BNST (ovBNST) of control mice to compare its effect on the anxiety/depressive-like behavior.**

(A): PKA-selective antagonist H89 (Control+H89; n=7) has no significant effect on the sucrose preference percentage of Control mice (n=15) (p>0.05).

(B): PKA-selective antagonist H89 (Control+H89; n=7) has no significant effect on the duration time that mice spent in the open arm of EPM test compared with Control mice (n=12) (p>0.05).

(C): H89 (Control+H89; n=7) had no significant effect on the frequency that mice entried into the open arm of EPM test compared with Control mice (Control; n=11) (p>0.05).

(D): H89 (Control +H89; n=7) had no significant effect on the distance that the Control mice (Control, n=13) (p>0.05) traveled in the center of open field (OF) test.

(E): H89 (Control +H89; n=6) had no significant effect on the duration that the Control mice (Control; n=13) (p>0.05) spent in the center of open field (OF) test.

(F): H89 (Control +H89; n=7) has no significant effect on the frequency that the Control mice (Control; n=13) (p>0.05) entried into the center of open field (OF) test.

(G): H89 (Control+H89; n=5) had no significant effect on the latency time of Control mice (Control; n=15) (p>0.05) to eat food pellet in the novelty suppressed feeding (NSF) test.

**Fig. S10**: **We propose an additional working model**, demonstrating that in which, under basal status, unphosphorylated (and active) STEP de-phosphorylates M-channel subunit and AMPA receptor (GluR1) subunit on the postsynaptic membrane, facilitating AMPAR (GluR1) internalization. After CVMS, activation of PKA will initiate STEP phosphorylation and its inactivation, which will in turn release a phosphorylation brake and consequently result in extra-phosphorylation of both membrane M-channel and surface GluR1 (and more GluR1 surface trafficking and expression). This will lead to a concomitantly decreased M-current and increased mEPSC amplitude. In this way, this indirect parallel pathway (through releasing phosphorylation brake by inactivating STEP) will converge with the direct pathway (by PKA phosphorylation) to mediate hyper- phosphorylation of M-channel and AMPA receptor GluR1 subunit seen in the chronic stress condition.
