## Supplementary figures and images for "Corticotropin-Releasing Hormone Signaling in the Oval Bed Nucleus of the Stria Terminalis Mediates Chronic Stress-Induced Negative Valence Behaviors Associated with Anxiety"

### Supplemental Figure 1

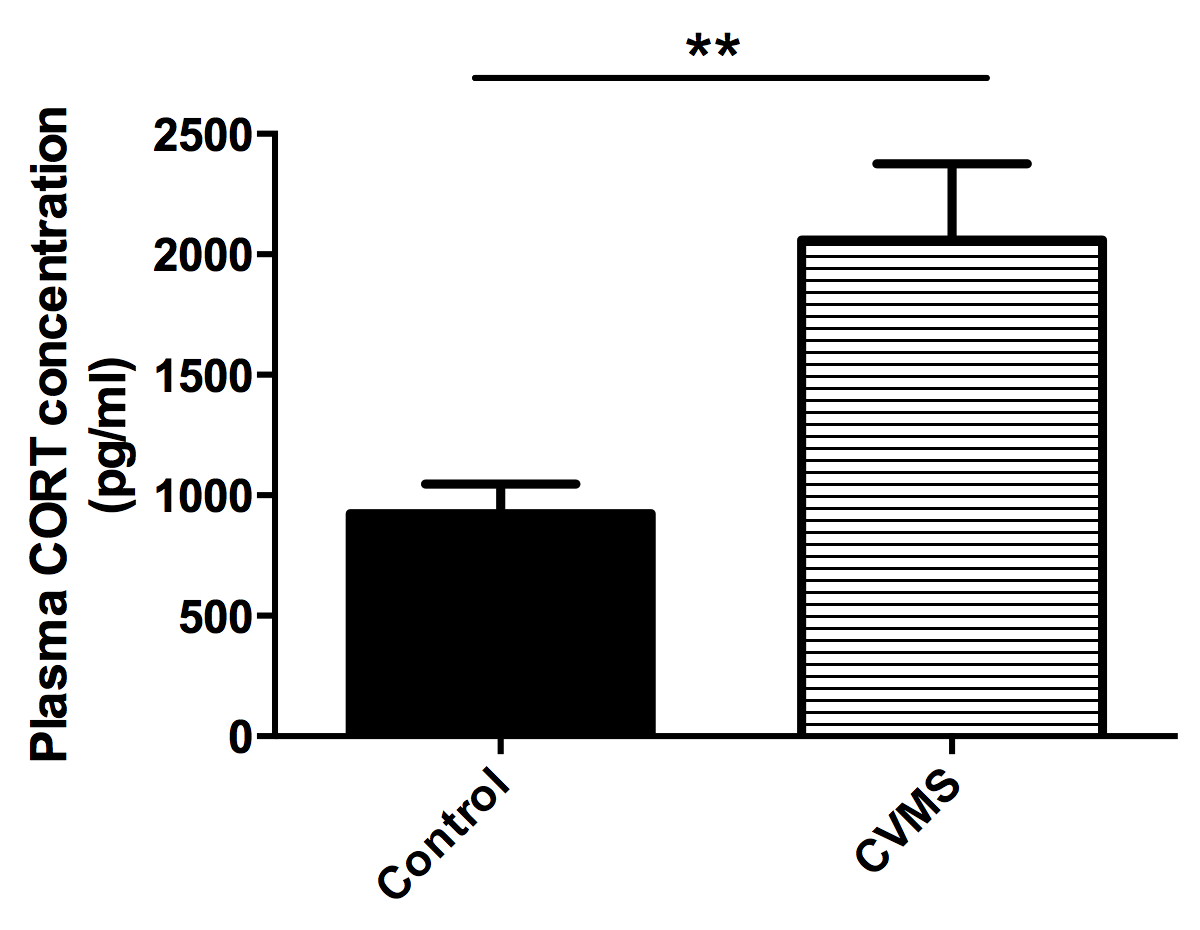

### Supplemental Figure 2

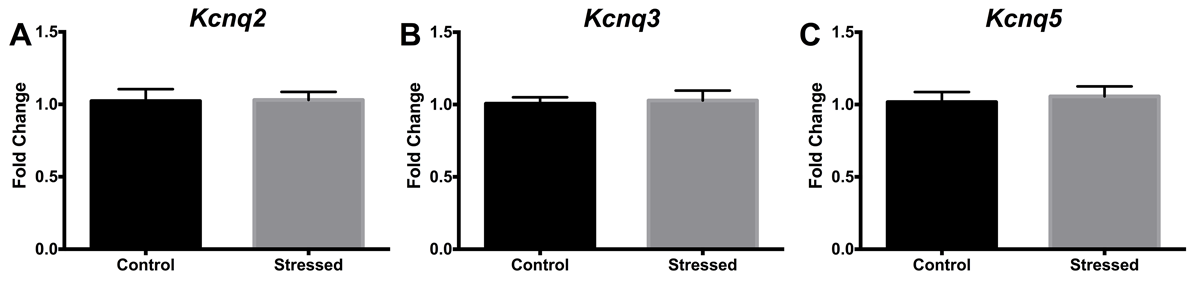

### Supplemental Figure 3

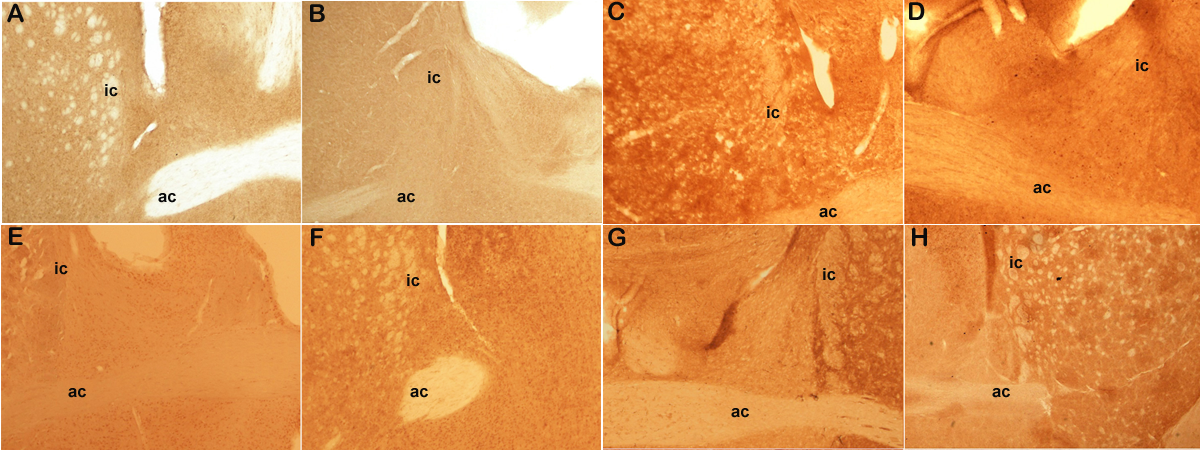

### Supplemental Figure 4 Part 1

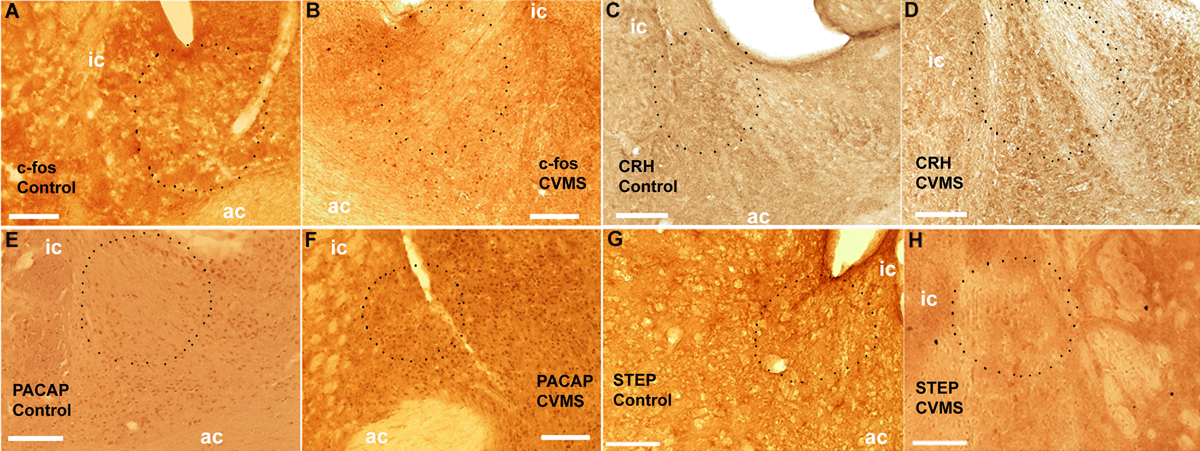

### Supplemental Figure 4 Part 2

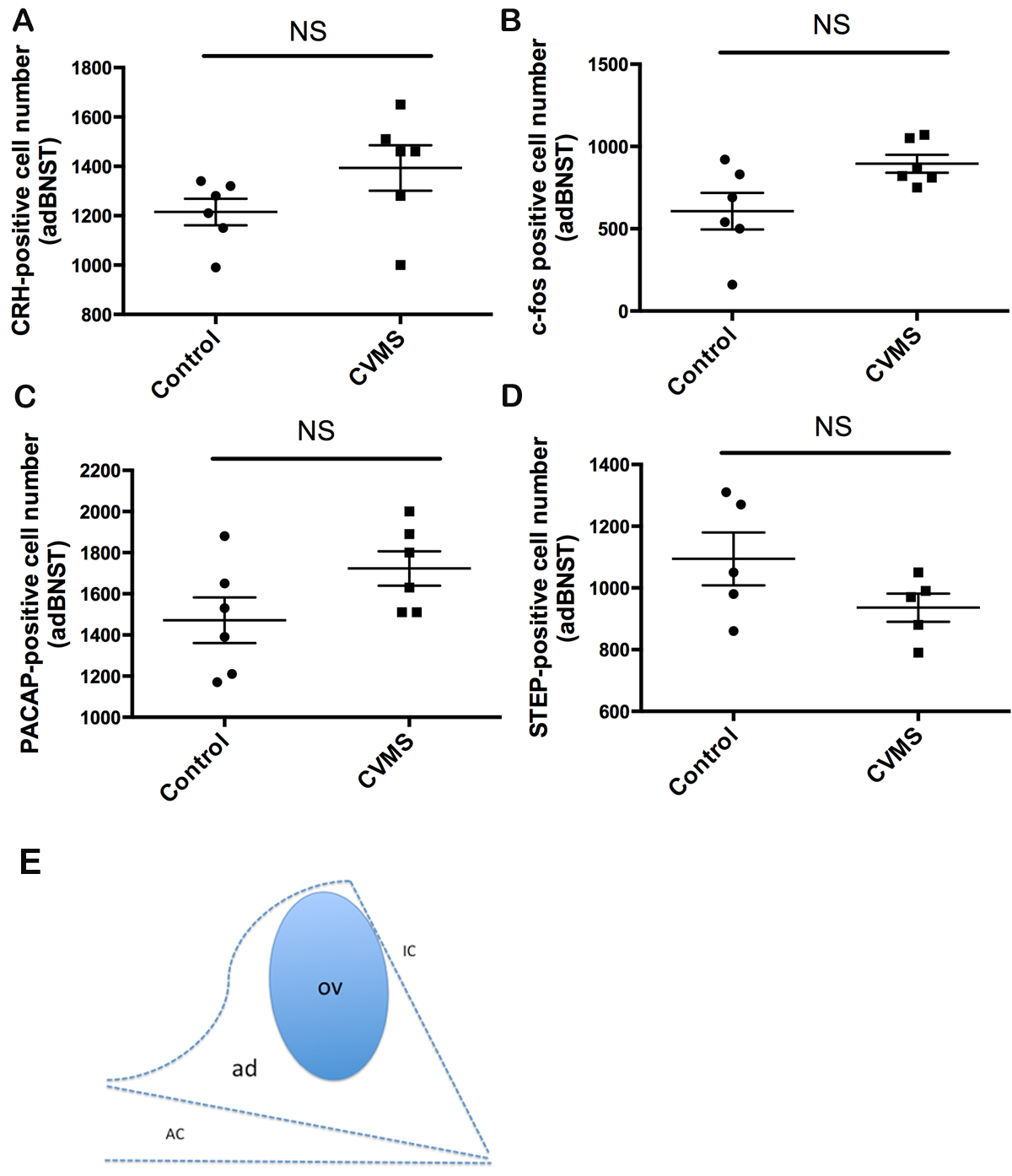

### Supplemental Figure 5

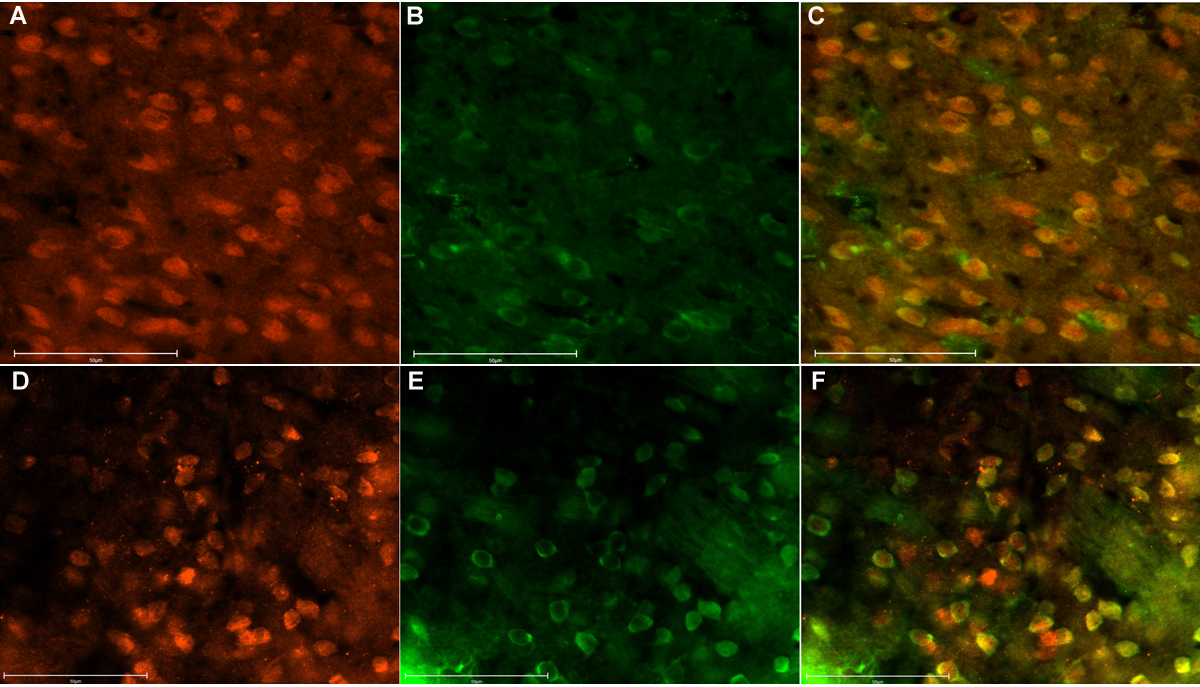

### Supplemental Figure 6

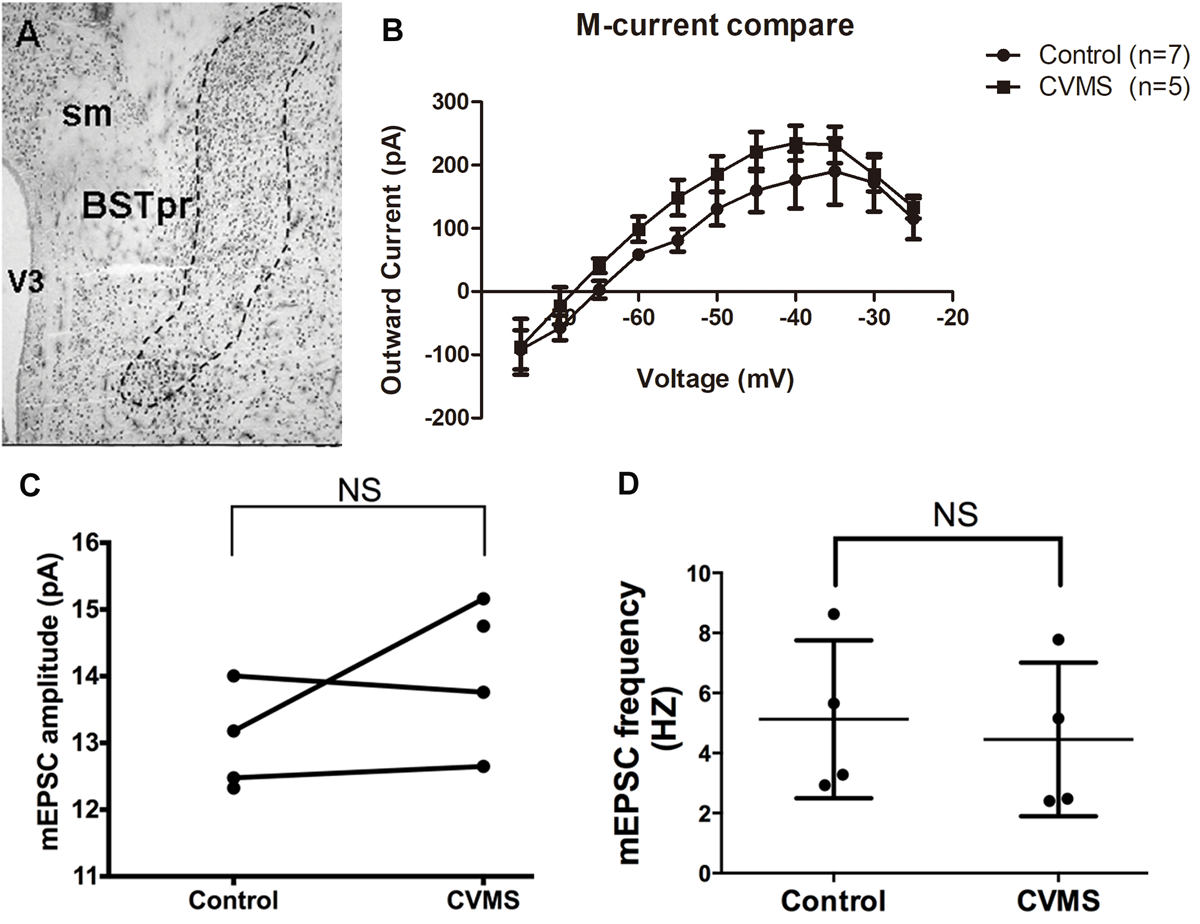

### Supplemental Figure 7 Part 1

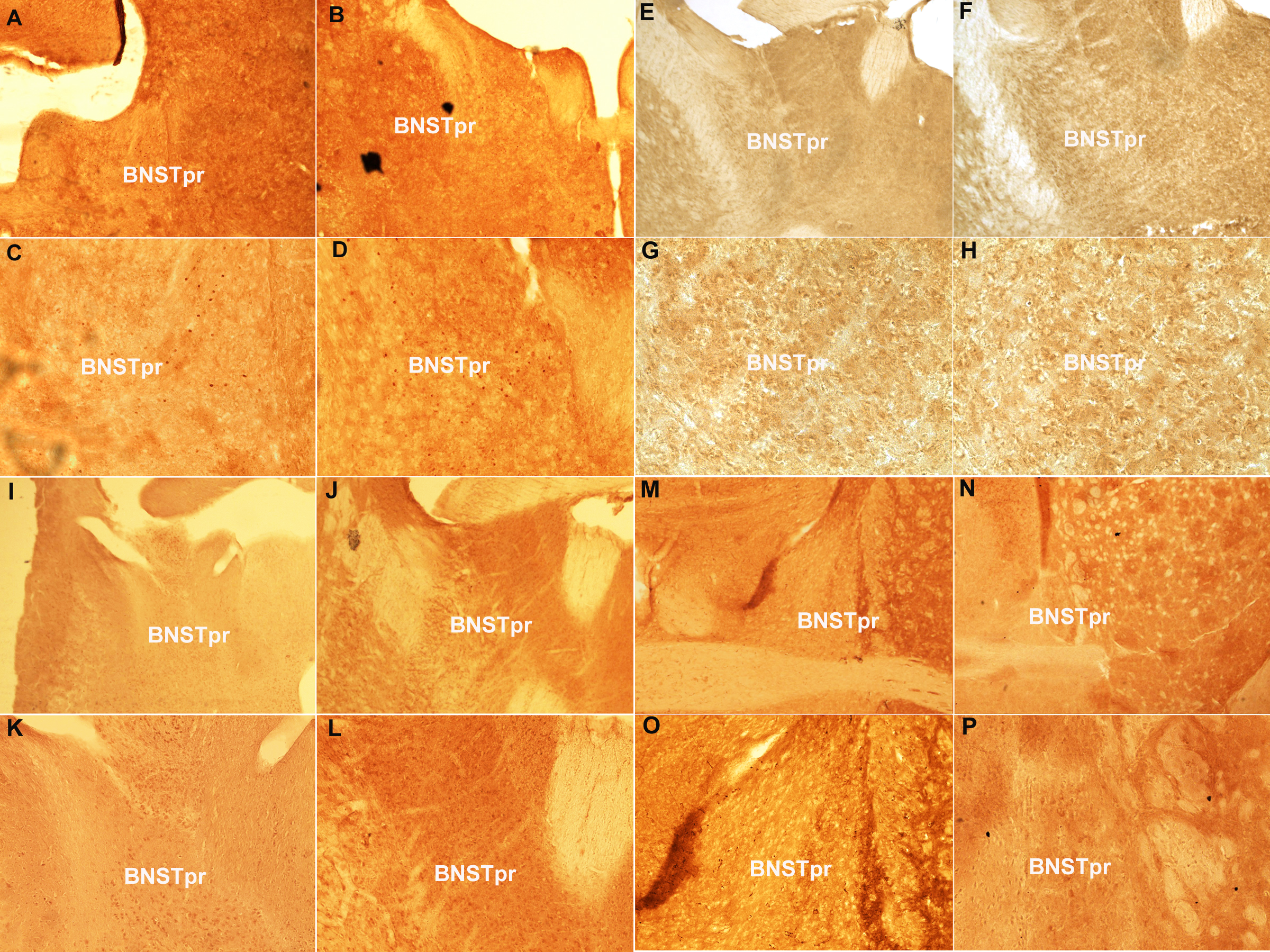

### Supplemental Figure 7 Part 2

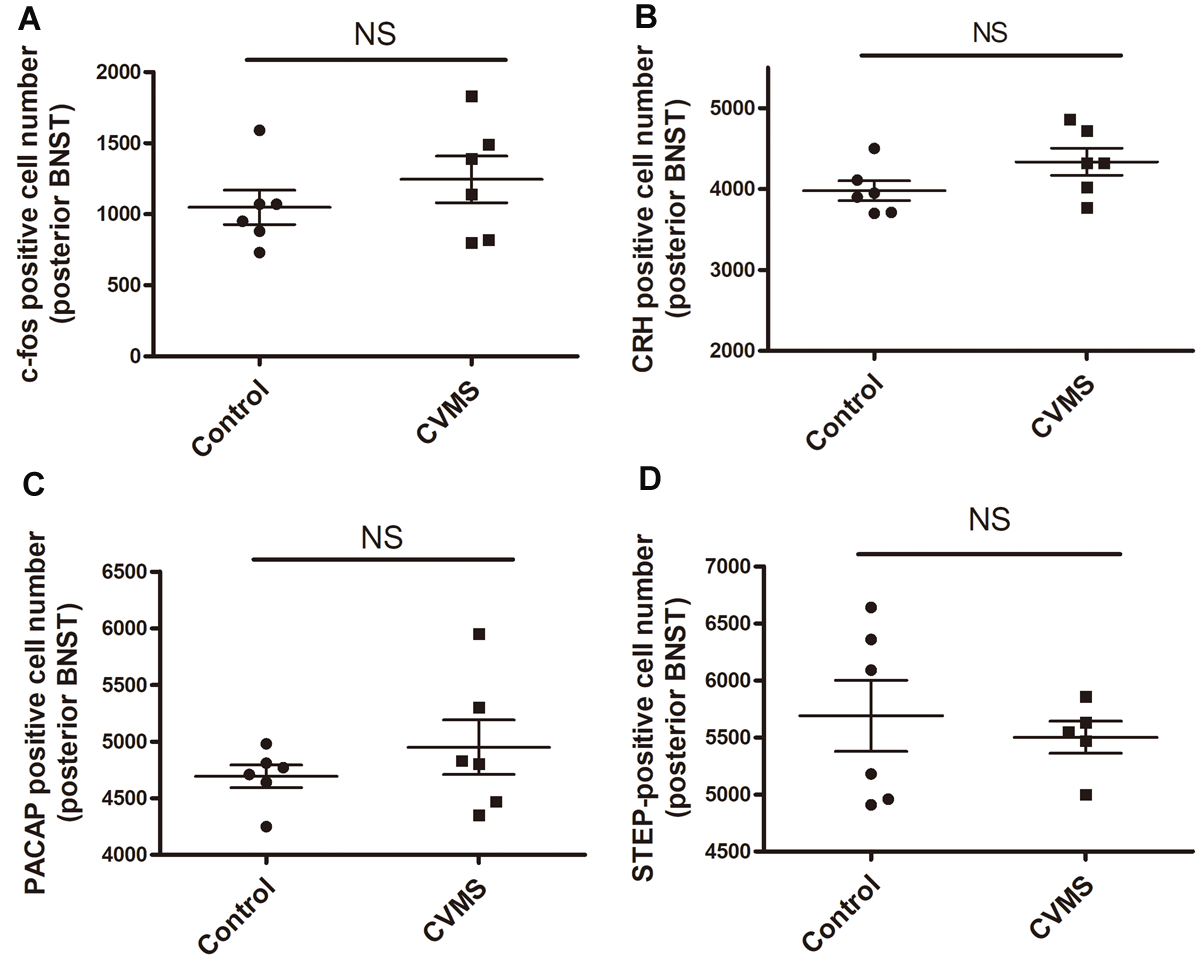

### Supplemental Figure 7 Part 3

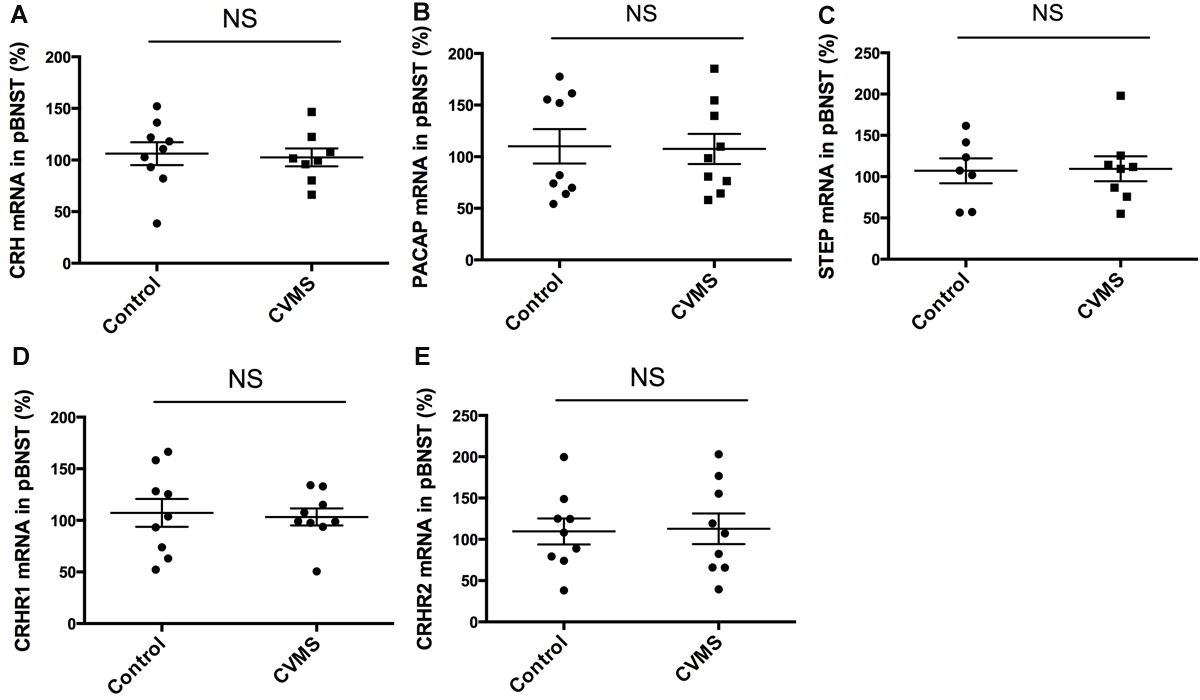

### Supplemental Figure 8

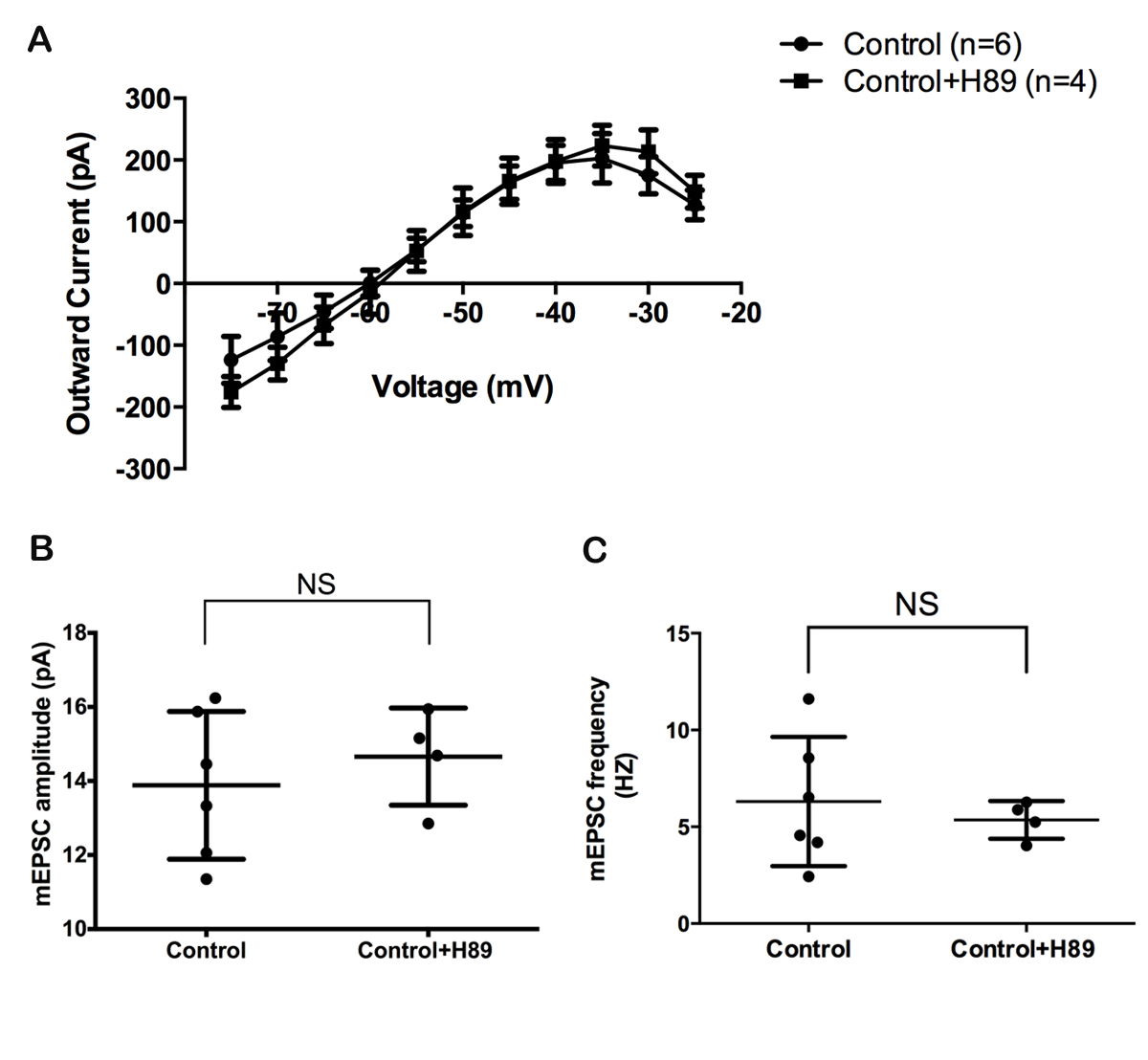

### Supplemental Figure 9

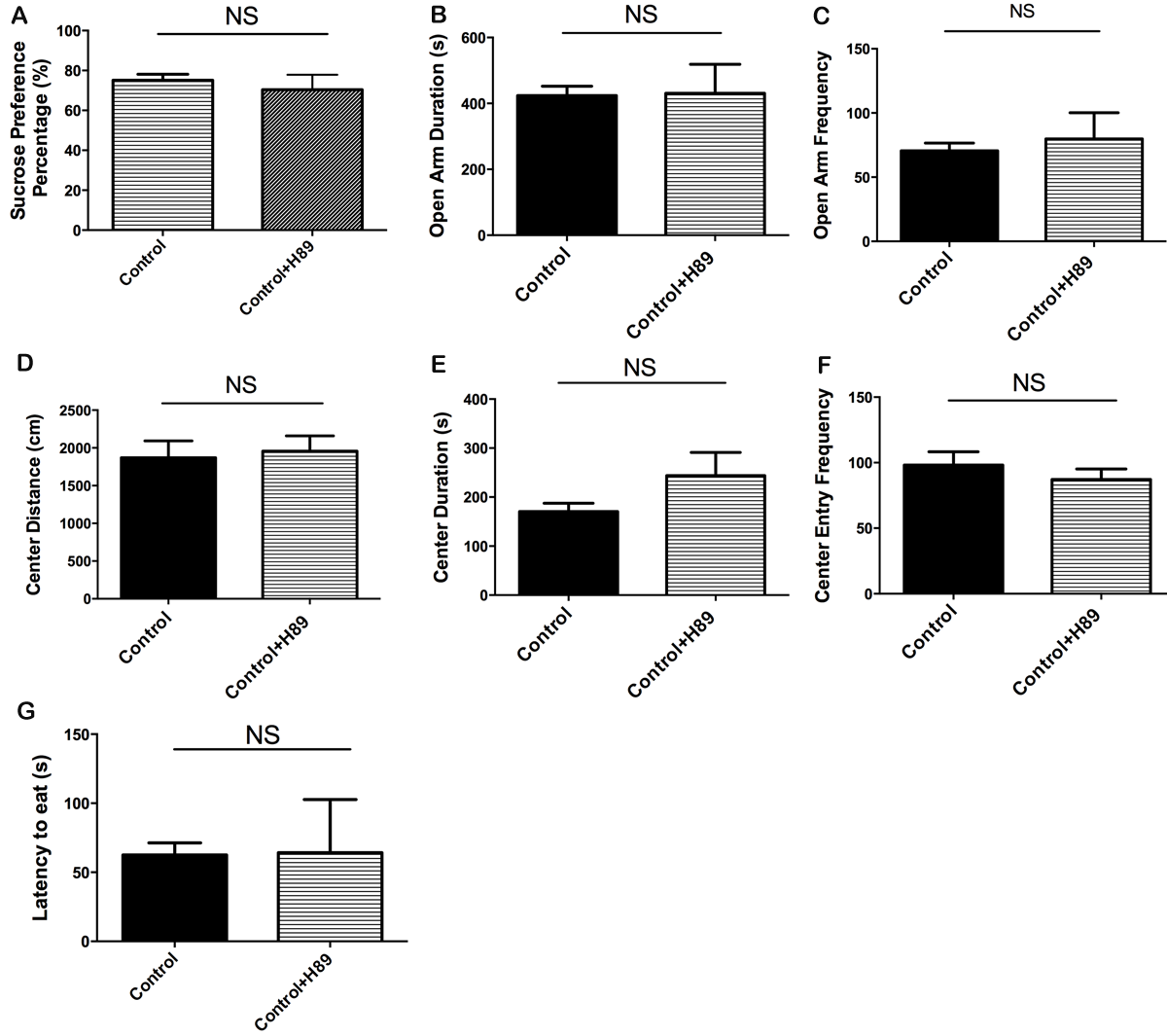

### Supplemental Figure 10

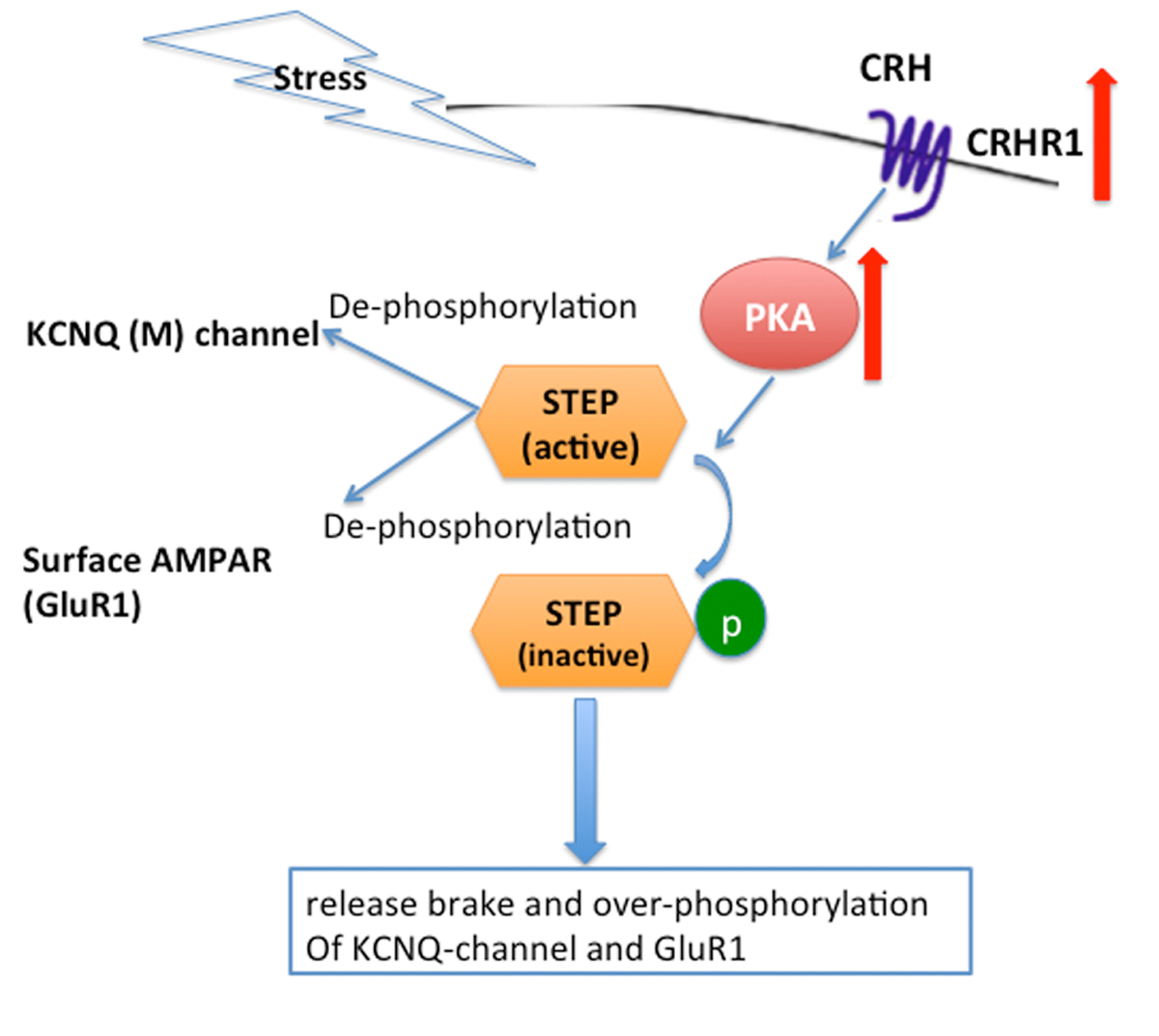
